# Mesenchymal *Vangl* facilitates airway elongation and widening independently of the planar cell polarity complex

**DOI:** 10.1101/2023.07.03.547543

**Authors:** Sarah V. Paramore, Katharine Goodwin, Danelle Devenport, Celeste M. Nelson

## Abstract

A hallmark of mammalian lungs is the fractal nature of the bronchial tree. In the adult, each successive generation of airways is a fraction of the size of the parental branch. This fractal structure is physiologically beneficial, as it minimizes the energy needed for breathing. Achieving this pattern likely requires precise control of airway length and diameter, as the branches of the embryonic airways initially lack the fractal scaling observed in those of the adult lung. In epithelial monolayers and tubes, directional growth can be regulated by the planar cell polarity (PCP) complex. Here, we comprehensively characterized the roles of PCP-complex components in airway initiation, elongation, and widening during branching morphogenesis of the murine lung. Using tissue-specific knockout mice, we surprisingly found that branching morphogenesis proceeds independently of PCP-component expression in the developing airway epithelium. Instead, we found a novel, *Celsr1*-independent role for the PCP component *Vangl* in the pulmonary mesenchyme. Specifically, mesenchymal loss of *Vangl1/2* leads to defects in branch initiation, elongation, and widening. At the cellular level, we observe changes in the shape of smooth muscle cells that indicate a potential defect in collective mesenchymal rearrangements, which we hypothesize are necessary for lung morphogenesis. Our data thus reveal an explicit function for *Vangl* that is independent of the core PCP complex, suggesting a functional diversification of PCP components in vertebrate development. These data also reveal an essential role for the embryonic mesenchyme in generating the fractal structure of airways of the mature lung.

## Introduction

The airway epithelium of the mammalian lung contains thousands of terminal ends that are generated via recursive rounds of stereotyped branching (Metzger et al., 2008). The epithelium can branch by initiating a bifurcation, wherein the tip of an existing branch folds in half, or a domain branch, wherein a daughter branch erupts from the side of an existing airway. These processes are influenced by the patterned differentiation of smooth muscle cells, which are derived from the surrounding mesenchyme and provide mechanical and chemical signals to the developing epithelium (Kim et al., 2015, Goodwin et al., 2019, Goodwin et al., 2022, Goodwin et al., 2023). After airway branches are initially established, the resulting tubes continue to increase in length and diameter as the lung develops, culminating in a fractal pattern in which the proximal airways are longer and wider than the distal airways (Tanabe et al., 2020, Nelson et al., 1990). This coordinated increase in airway size is essential for proper flow of air after birth.

Although branch initiation has been investigated extensively, the mechanisms that drive epithelial lengthening and widening to generate the fractal pattern of airways remain poorly understood. In the kidney, tubule elongation is driven by epithelial-intrinsic mechanisms, such as oriented cell divisions and convergent extension. Oriented cell divisions have likewise been observed during branching morphogenesis of the lung (Tang et al., 2018, Tang et al., 2011), where they were also found to occur predominately in the axial direction, promoting airway lengthening. Given that airways also widen in diameter over developmental time, epithelial remodeling in the developing lung is unlikely to result solely from axially oriented cell divisions. To achieve its final fractal architecture, the embryonic airway must deploy additional mechanisms to tune its local length and diameter.

Planar cell polarity (PCP) refers to the long-range, collective polarization of cells along a tissue plane and is controlled by a highly conserved molecular module known as the core PCP complex. First characterized in epithelia, the core PCP complex consists of a set of transmembrane and cytoplasmic proteins that localize asymmetrically at intercellular junctions (Devenport, 2014). The transmembrane protein Vangl complexes with the cytoplasmic protein Prickle and localizes to the opposite side of the cell as the complex formed by transmembrane Frizzled (Fzd) and cytoplasmic Dishevelled. Celsr, an atypical cadherin, localizes to both sides of intercellular junctions and connects Fzd and Vangl complexes between neighboring cells through homotypic adhesion (Devenport, 2014). Of the Fzd receptors in mice, Fzd3 and Fzd6 function in the PCP complex (Wang et al., 2006). Broadly, PCP serves as a compass that can direct individual cell behaviors, such as oriented cell divisions, or collective cell behaviors, such as convergent extension (Tada and Smith, 2000, Heisenberg et al., 2000, Wallingford et al., 2000). Loss of a functional PCP complex leads to severe developmental defects in many organ systems, including a failure to close the neural tube (Curtin et al., 2003, Torban et al., 2004, Wang et al., 2006). Thus, PCP is an essential complex that regulates the shape of tissues and organs across development.

Although best understood in simple cellular sheets and tubes, PCP has also been implicated in the development of branched organs, including the kidney and mammary gland. During morphogenesis of the kidney, PCP drives cell intercalation through the formation and resolution of multicellular rosettes leading to convergent extension and thus elongation of kidney tubules; when PCP is lost, kidney tubules remain short and wide (Lienkamp et al., 2012, Kunimoto et al., 2017). During morphogenesis of the mammary gland, loss of the core PCP protein Vangl2 also results in dysregulation of tube size; however, the cellular basis of the phenotype is unknown (Smith et al., 2019). Two core transmembrane PCP components, *Vangl2* and *Celsr1*, are essential for aspects of lung development and homeostasis. In the upper airways, PCP orients the coordinated beating of ciliated cells that line the trachea (Vladar et al., 2012). Cilia are misoriented in PCP mutants and fail to polarize along the proximal-distal axis of the lung epithelium. The PCP pathway has also been implicated in branching morphogenesis, as PCP mutants display alterations in the number (Yates et al., 2010) and positioning (Zhang et al., 2022) of branches as compared to littermate controls. However, the cellular mechanisms by which loss of PCP affects morphogenesis of the embryonic lung have not been reported. Given that PCP proteins are expressed in the lung from the onset of branching (Vladar et al., 2012), it is possible that PCP influences the fractal shape of the airways by regulating polarized cellular behaviors throughout branch initiation, elongation, and widening. Here, we investigate the links between the PCP complex and formation of the fractal tree of airways during lung development. We reveal that *Celsr1* is expressed only in the lung epithelium and mesothelium, while *Vangl2* is expressed ubiquitously throughout the tissues of the lung, including in the pulmonary mesenchyme. We find that *Vangl2-*mutant lungs, but not *Celsr1-* mutant lungs, exhibit a decrease in branch number. However, *Vangl2*-mutant lungs still form branches in the expected stereotyped pattern. To define the tissue-specific role of *Vangl* in airway branching morphogenesis, we take advantage of the *Cre-Lox* system to specifically delete *Vangl1/2* from the airway epithelium or the pulmonary mesenchyme. We show that loss of epithelial *Vangl1/2*, and thus loss of epithelial PCP, does not affect branch initiation, elongation, or widening. Surprisingly, we find a significant role for *Vangl1/2* in the pulmonary mesenchyme: loss of mesenchymal *Vangl1/2* leads to a severe reduction in branch initiation and striking defects in airway elongation and widening. Our data reveal that the pulmonary mesenchyme plays a key role in shaping the morphology of the branching airway epithelium in a manner dependent on *Vangl*, but independent of the core PCP complex.

## Results

### Vangl2-mutant lungs have a reduction in airway epithelial volume and branch number

To examine the expression of PCP components (**Fig. 1A**) during lung development, we harvested lungs from embryonic stages spanning the days of epithelial branching morphogenesis (*E*11, *E*13, *E*15) and used immunofluorescence analysis to reveal the localization of PCP proteins. We found that throughout the branching process, Vangl2 is membrane-localized in epithelial and mesenchymal cells (**Fig. 1B, Supplemental Fig. 1A**), whereas Celsr1 localizes to the membrane only in epithelial cells (**Fig. 1C, Supplemental Fig. 1B**). Fzd6 is localized to the membranes of both epithelial and endothelial cells (**Fig. 1D, Supplemental Fig. 1C**). Consistently, single-cell RNA-sequencing (scRNA-seq) analysis of *E*11.5 lungs (Goodwin et al., 2022) revealed that while *Vangl2* is expressed predominantly in the airway epithelium, undifferentiated mesenchyme, and mesothelium (**Fig. 1E**), *Celsr1* is only expressed in the epithelium and mesothelium (**Fig. 1F**). *Fzd6* expression was relatively low in our dataset, likely due to underreading of transcript (**Fig. 1G**).

**Figure 1.**
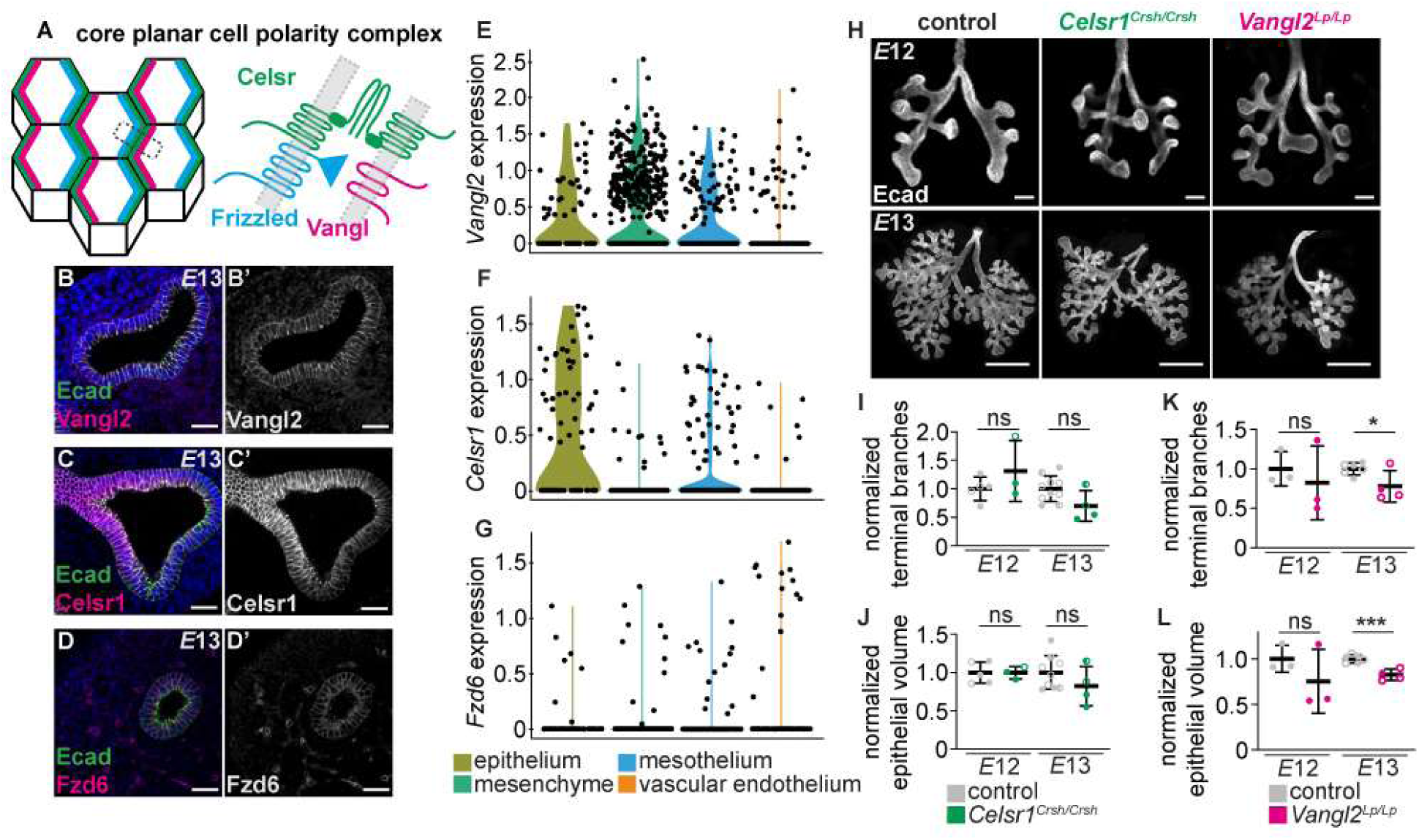
Airway branching morphogenesis is slightly delayed in *Vangl2^Lp/Lp^* mutants. **A,** Schematic illustrating the core PCP complex. **B-D,** Sections (10-µm-thick) from *E*13 lungs immunostained for Vangl2, Celsr1, or Fzd6 (magenta) and E-cadherin (Ecad; green). Nuclei are counterstained with Hoechst (blue); scale bars, 25 µm. **E-G,** Violin plots showing transcript levels of *Vangl2*, *Celsr1,* and *Fzd6* from scRNA-sequencing of *E*11.5-12 lungs. **H,** Z-projections of confocal slices acquired from cleared, whole-mount lungs from *E*12 and *E*13 embryos immunostained for Ecad; scale bars, 500 µm. **I,** Normalized number of terminal branches in control and *Celsr1^Crsh/Crsh^* lungs at *E*12 (*n*=5 control and *n*=3 mutant lungs, *p*=0.2678 via unpaired Student’s t-test) and *E*13 (*n*=10 control and *n*=4 mutant lungs, *p*=0.0541 via unpaired Student’s t-test). **J,** Normalized epithelial volume in control and *Celsr1^Crsh/Crsh^* lungs at *E*12 (*n*=5 control and *n*=3 mutant lungs, *p*=0.9887 via unpaired Student’s t-test) and *E*13 (n=9 control and *n*=4 mutant lungs, *p*=0.2277 via unpaired Student’s t-test). **K,** Normalized number of terminal branches in control and *Vangl2^Lp/Lp^* lungs at *E*12 (*n*=3 control and *n*=3 mutant lungs, *p*=0.5850 via unpaired Student’s t-test) and *E*13 (*n*=6 control and *n*=4 mutant lungs, *p*=0.0360 via unpaired Student’s t-test). **L,** Normalized epithelial volume in control and *Vangl2^Lp/Lp^* lungs at *E*12 (*n*=3 control and *n*=3 mutant lungs, *p*=0.3259 via unpaired Student’s t-test) and *E*13 (*n*=6 control and *n*=4 mutant lungs, *p*=0.0007 via unpaired Student’s t-test). In all graphs, different shapes represent distinct experimental replicates; shown are mean ± s.d; * p < 0.05; *** p < 0.001.

To begin to define the role of PCP in lung development, we harvested lungs from *Celsr1^Crsh/Crsh^* and *Vangl2^Lp/Lp^* embryos, both of which harbor dominant-negative mutations that disrupt the asymmetric localization of PCP components as well as PCP function (Devenport and Fuchs, 2008, Curtin et al., 2003, Stahley et al., 2021, Yin et al., 2012, Vladar et al., 2012). Specifically, the *Celsr1^Crsh^* allele harbors a single amino acid substitution between two cadherin repeats in the extracellular domain of the protein, thus affecting cis-interactions and assembly of PCP complexes (Stahley et al., 2021). Similarly, the *Vangl2^Lp^*mutation blocks PCP complex assembly due to a point mutation in its ER export sequence that disrupts both Vangl2 and Vangl1 trafficking to the cell surface (Yin et al., 2012). At *E*12 and *E*13, *Celsr1^Crsh/Crsh^* lungs form the expected pattern of branches with no significant differences from littermate controls (**Fig. 1H-J, Supplemental Fig. 1D-E**). Consistently, we found that when cultured for two days *ex vivo*, *Celsr1^Crsh/Crsh^* lung explants branch at the same rate as controls, yielding a similar number of terminal branches with a comparable fold change in number of terminal branches (**Supplemental Fig. 1F-H**). *Celsr1* is therefore surprisingly dispensable for branch initiation in the airway epithelium.

In contrast, *E*13 *Vangl2^Lp/Lp^* lungs are significantly smaller and display a significant reduction in the number of terminal branches and overall epithelial volume compared to lungs of littermate controls (**Fig. 1H, 1K-L, Supplemental Fig. 1I-J**). The striking difference in airway architecture between *Celsr1*-mutant and *Vangl2*-mutant lungs led us to hypothesize that *Vangl2* may function independently of *Celsr1* in airway epithelial morphogenesis.

### Removal of Vangl1/2 from the lung epithelium does not affect branch initiation

Because *Vangl1/2* is expressed ubiquitously in the lung, we generated tissue-specific conditional *Vangl1/2* knockout mice to separately assess *Vangl1/2* function in the epithelium and pulmonary mesenchyme. We first generated *ShhCre; Vangl1 ^fl/fl^; Vangl2 ^fl/fl^*(epithelial-conditional-knockout or epiCKO) animals. In contrast to the conclusions of previous work (Zhang et al., 2022), we found that the *ShhCre* driver efficiently deletes *Vangl1/2*, as we observed loss of detectable Vangl2 protein in the epithelium, but not the mesenchyme, of epiCKO lungs (**Supplemental Fig. 2A**). Further, we found that removal of *Vangl1/2* in the epiCKO animals leads to a loss of Celsr1 polarization in the trachea (Paramore et al., 2022); thus, using the *ShhCre* driver to delete *Vangl1/2* causes loss of a functional PCP complex in the embryonic lung epithelium.

Whereas epiCKO lungs are indistinguishable from controls at *E*12, mutant lungs show a significant reduction in the number of terminal branches by *E*13 (**Fig. 2A-B, Supplemental Fig. 2B**). However, we also found that epiCKO embryos are significantly smaller than their littermate controls and appear to be developmentally delayed, likely due to loss of *Vangl1/2* in *Shh*-expressing tissues outside of the lung (**Fig. 2C-D**). It was therefore possible that the decrease in branch number observed in epiCKO lungs was due to the smaller size and developmental delay of the embryo itself, rather than due to an intrinsic requirement for *Vangl1/2* in the lung epithelium.

**Figure 2.**
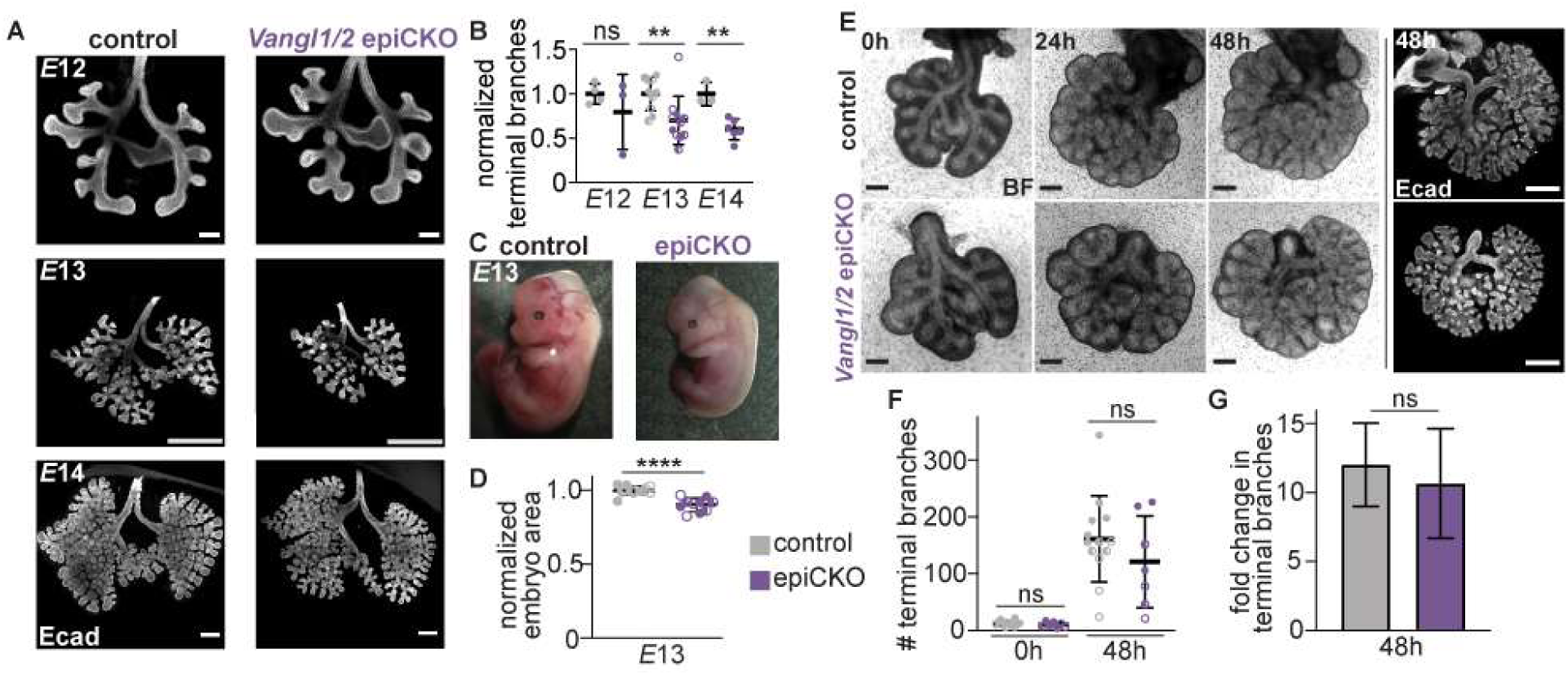
Epithelial *Vangl* is not required for branch initiation. **A,** Z-projection of confocal slices acquired from cleared, whole-mount *E*12-14 lungs from control and epiCKO embryos. **B,** Normalized number of terminal branches in control and epiCKO lungs at *E*12 (*n*=4 control and *n*=3 mutant lungs, *p*=0.3883 via unpaired Student’s t-test), *E*13 (*n*=10 control and *n*=11 mutant lungs, *p*=0.0090 via unpaired Student’s t-test), and E14 (*n*=3 control and *n*=6 mutant lungs, *p*=0.0023 via unpaired Student’s t-test). **C,** Representative images of *E*13 control and epiCKO embryos. **D,** Quantification of normalized projected areas of *E*13 control and epiCKO embryos (*n*=11 control and *n*=10 mutant embryos, *p*=0.0006 via unpaired Student’s t-test). **E,** Representative time-lapse images of control and epiCKO lungs dissected at *E*12 and cultured for 48 h; scale bars, 250 µm. **F,** Number of terminal branches at 0 h and 48 h of culture for control and epiCKO lungs (*n*=13 control and *n*=7 mutants, *p*=0.2257 at 0 h, *p*=0.2845 at 48 h via unpaired Student’s t-test). **G,** Fold change in number of terminal branches after 48 h of culture for control and epiCKO lungs (*n*=13 control and *n*=7 mutants, *p*=0.4008 via unpaired Student’s t-test). Shown are mean ± s.d; ** p < 0.01; *** p < 0.001; **** p < 0.0001. In all graphs, different shapes represent distinct experimental replicates.

To distinguish between these two possibilities, we tested whether epiCKO lungs retain the same capacity for branching as control lungs when removed from the constraints of the embryonic chest cavity. We dissected lungs from *E*12 embryos, a stage at which we did not observe a difference between mutants and controls, and monitored their rate of epithelial branching for 48 h in culture. We found that when cultured *ex vivo*, epiCKO lungs branch at the same rate as controls (**Fig. 2E-G, Supplemental Fig. 2C**). It is therefore likely that any delay in branching observed in epiCKO lungs results from the developmental delay of the embryo, rather than from loss of epithelial PCP per se. Because the number of branches is unaffected by loss of either *Celsr1* or epithelial *Vangl1/2*, we conclude that branch initiation occurs independently of the core PCP complex in the murine lung.

### Removal of Vangl1/2 from the lung mesenchyme leads to defects in branch initiation

In addition to being expressed in the lung epithelium, the core PCP component *Vangl2* is also expressed at high levels in the lung mesenchyme (**Fig. 1B, E**). Given that epiCKO lungs initiate branches normally, we hypothesized that the branching phenotype observed in *Vangl2^Lp/Lp^* lungs might indicate a role for *Vangl* in the pulmonary mesenchyme. To test this hypothesis, we began by breeding a mesenchymal *Vangl1/2* knockout: a *Dermo1Cre; Vangl1 ^fl/fl^; Vangl2^fl/fl^* mouse line, in which *Cre* is expressed broadly in the splanchnic mesoderm, including in the pulmonary mesenchyme, but is absent from the airway epithelium (Chen et al., 2008). Notably, the *Dermo1Cre* allele and *Vangl2* gene are on the same chromosome; thus, generating homozygous embryos necessitates a meiotic crossover event. In contrast to the conclusions made in previous work (Zhang et al., 2020), we found that *Dermo1Cre; Vangl1 ^fl/fl^; Vangl2^fl/fl^* embryos survive early development, as we could recover homozygotes as late as *E*18.5. Surprisingly, lungs from *E*13.5 mutant embryos appear phenotypically normal and do not recapitulate the defects in branching observed in the *Vangl2^Lp/Lp^* lungs (**Supplemental Fig. 3A-B**). Immunostaining for Vangl2 in control and *Dermo1Cre; Vangl1 ^fl/fl^; Vangl2^fl/fl^* lungs revealed significant levels of Vangl2 protein in the mesenchyme at *E*13 (**Supplemental Fig. 3C**). Thus, we conclude that either *Dermo1Cre* does not efficiently delete *Vangl2* during these stages of development, or that Vangl2 protein is stable and persists in the mesenchyme long after Cre-mediated recombination.

To circumvent this issue, we generated a *Tbx4-rtTA; Tet-O-Cre; Vangl1^fl/fl^; Vangl2^fl/fl^* (inducible-mesenchymal-conditional-knockout or inducible-mesCKO) mouse line, in which expression of *Cre* is restricted specifically to the pulmonary mesenchyme (Zhang et al., 2013). In contrast to the *Dermo1Cre* driver, we found that the *Tbx4-rtTA; Tet-O-Cre* driver results in significant loss of detectable Vangl2 protein in the mesenchyme by *E*12.5 when doxycycline-containing drinking water is introduced to the dam at *E*8. (**Supplemental Fig. 4A**). To ensure full deletion of the desired alleles during the pseudoglandular stage of lung development, we proceeded to use the *Tbx4-rtTA; Tet-O-Cre* system.

We analyzed lungs from these inducible-mesCKO embryos at *E*12-14. By *E*13, lungs of inducible-mesCKO embryos are noticeably smaller than those of their littermate controls (**Fig. 3A-B, Supplemental Fig. 4B**). In contrast to epiCKO lungs, this reduction in size cannot be attributed to an embryo-wide developmental delay: inducible-mesCKO embryos are the same size as control embryos (**Fig. 3C-D**). Consistent with their smaller size, inducible-mesCKO lungs show a significant reduction in the number of terminal branches at *E*14 (**Fig. 3E**). The new branches that do form appear to be correctly stereotyped and shaped, suggesting that branch initiation might be delayed but morphologically normal in mutants as compared to controls. Consistently, *E*12.5 inducible-mesCKO lungs form fewer branches when cultured *ex vivo* as compared to control lungs (**Fig. 4F-H**). Thus, deletion of *Vangl1/2* from the pulmonary mesenchyme leads to a reduction or delay in branch initiation.

**Figure 3.**
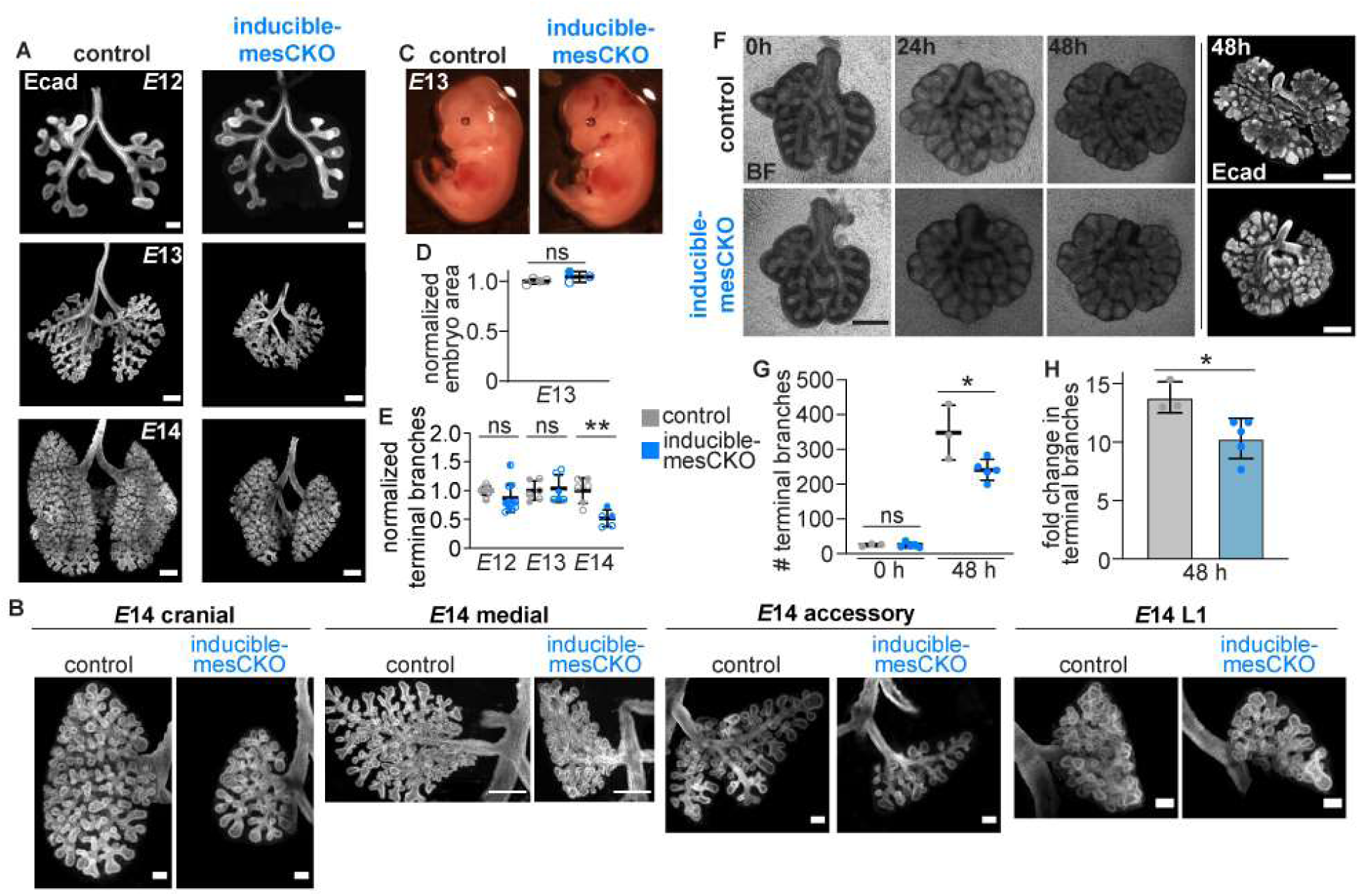
Loss of mesenchymal *Vangl* leads to defects in branch initiation. **A,** Z-projection of confocal slices acquired from cleared, whole-mount *E*12 (scale bar, 100 µm), *E*13 (scale bar, 250 µm), and *E*14 (scale bar, 250 µm) lungs from control and inducible-mesCKO embryos. **B,** Z-projection of confocal slides of individual *E*14 control and inducible-mesCKO cranial, medial, and accessory bronchi and L1 branch. **C,** Representative images of *E*13 control and inducible-mesCKO embryos. **D,** Quantification of normalized projected areas of *E*13 control and inducible-mesCKO embryos (*n*=4 control and *n*=3 mutant embryos, *p*=0.2077 via unpaired Student’s t-test). **E,** Normalized number of terminal branches in control and inducible-mesCKO lungs at *E*12 (*n*=8 control and *n*=11 mutant lungs, *p=*0.1550 via unpaired Student’s t-test), *E*13 (*n*=6 control and *n*=6 mutant lungs, *p=*0.7718 via unpaired Student’s t-test), and *E*14 (*n*=6 control and *n*=5 mutant lungs, *p=*0.0027 via unpaired Student’s t-test). **F,** Representative time-lapse images of control and inducible-mesCKO lungs dissected at *E*12 and cultured for 48 h; scale bars, 250 µm. **G,** Number of terminal branches at 0 h and 48 h of culture for control and inducible-mesCKO lungs (*n*=3 control and *n*=5 mutants, *p*=0.9060 at 0 h, *p*=0.0301 at 48 h via unpaired Student’s t-test). **H,** Fold change in number of terminal branches after 48 h of culture for control and inducible-mesCKO lungs (*n*=3 control and *n*=5 mutants, *p*=0.0239 via unpaired Student’s t-test).

**Figure 4.**
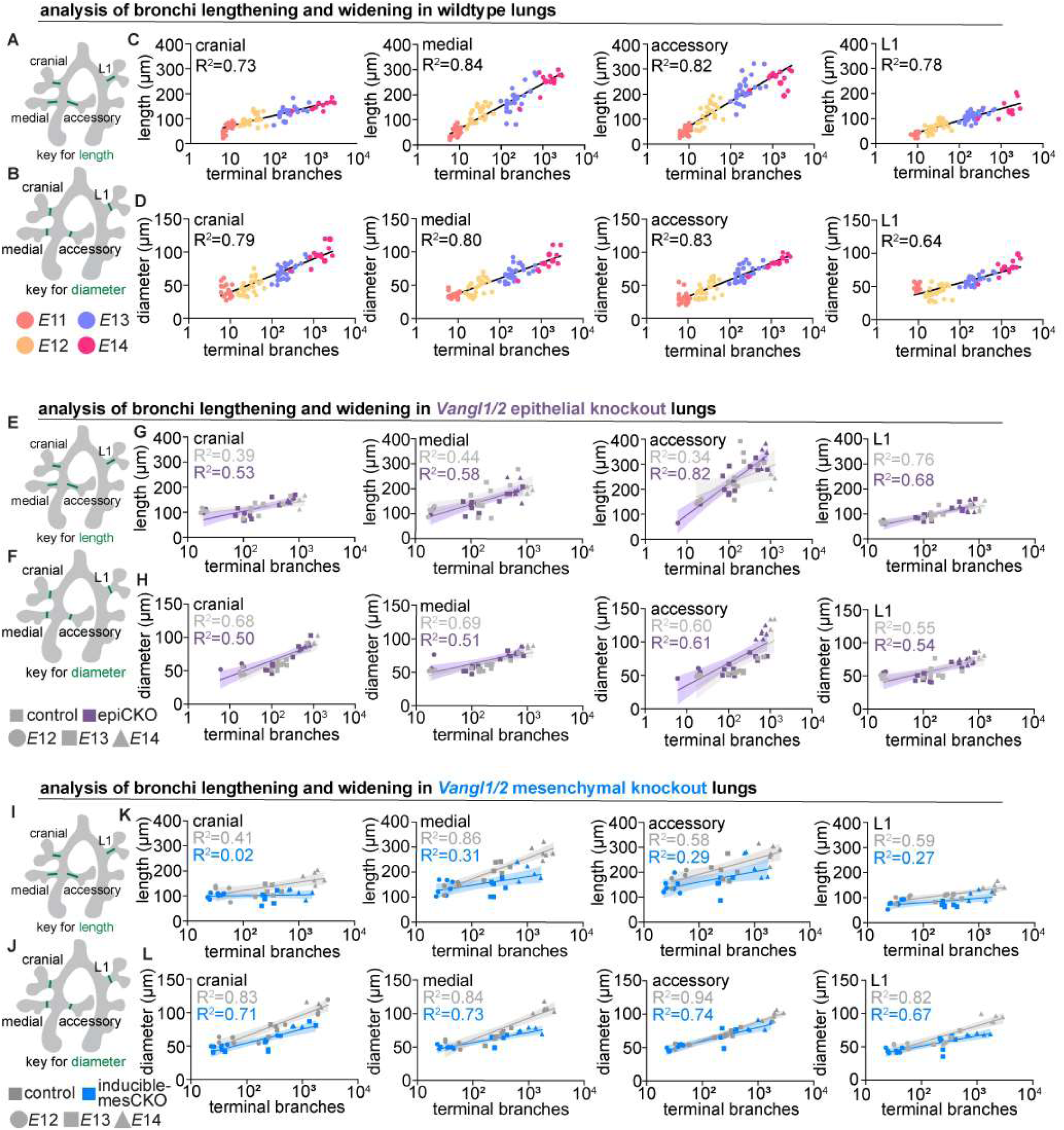
Branch elongation and widening both scale with branch complexity and require *Vangl* in the mesenchyme but not the epithelium. **A-B**, Schematics of lung with landmarks used for quantification of bronchus length and diameter. **C-D,** Quantification of bronchus length or diameter vs. number of terminal branches in *E*12-14 cranial, medial, and accessory bronchi and L1 branch from wild-type lungs. Shown are best-fit curves. **E-F,** Schematics of lung with landmarks used for quantification of bronchus length and diameter and key for epiCKO lungs. **G-H,** Quantification of bronchus length or diameter vs. number of terminal branches in *E*12-14 cranial, medial, and accessory bronchi and L1 branch from control and epiCKO lungs. Shown are best-fit curves and 95% confidence intervals. **I-J,** Schematics of lung with landmarks used for quantification of bronchus length and diameter and key for inducible-mesCKO lungs. **K-L,** Quantification of bronchus length or diameter vs. number of terminal branches in *E*12-14 cranial, medial, and accessory bronchi and L1 branch from control and inducible-mesCKO lungs. In all graphs, each dot represents one lung.

### Stereotyped scaling of branch length and diameter requires Vangl1/2 in the mesenchyme but not the epithelium

Though the stereotyped pattern of branch initiation has been described in detail (Metzger et al., 2008), less is known about the dynamics with which the airways elongate and widen to generate the fractal tree observed in the mature organ. Specifically, it is unclear how airway length and width scale as the lung grows. To assess changes in airway geometry over developmental time, we measured the lengths and diameters of four major airways: the bronchi of the cranial, medial, and accessory lobes, and the first branch on the left lobe (L1) (**Fig. 4A-D**). We selected these airways because they form during the earliest stages of branching, have clear morphological landmarks, and can be measured unambiguously in each lung. We plotted these measurements as a function of the number of terminal branches, as a proxy for developmental time. Surprisingly, we found that the four bronchi elongate at different rates. For example, the cranial bronchus increases in length ∼3-fold from *E*11 to *E*14.5, while the accessory bronchus increases in length ∼6-fold (**Fig. 4C**). In contrast, all four bronchi widen at similar rates, increasing ∼4-fold in diameter from *E*11 to *E*14.5 (**Fig. 4D**). These observations suggest that the relative growth of the airways, which generates the fractal structure of the epithelial tree, is spatially regulated during lung development.

The processes of airway lengthening and widening have been attributed primarily to behaviors by the epithelium itself. Specifically, airway epithelial cells have been reported to divide predominantly along the proximal-distal axis of the airway, which would lead to its lengthening (Tang et al., 2018). In organs such as the kidney, the core PCP complex directs oriented cell divisions and other cellular behaviors that promote lengthening, such as convergent extension (Kunimoto et al., 2017). Thus, though we found that the PCP complex was dispensable for branch initiation in the developing lung, we hypothesized that epithelial PCP may regulate airway lengthening and/or widening. To determine whether PCP is required in the epithelium for airway remodeling, we compared the lengths and diameters of control and epiCKO bronchi from *E*12-14.5. After generating best-fit curves for both the control and mutant data sets, we found that the 95% confidence intervals of our control and mutant data overlap for each bronchus analyzed (**Fig. 4E-H**). While the epiCKO embryos experience a developmental delay, the length and diameter of their bronchi still scale with the number of terminal branches in a manner comparable to control lungs. We therefore conclude that PCP in the epithelium is also dispensable for elongation and widening of the airways during the branching morphogenesis stages of lung development.

Because inducible-mesCKO lungs appear significantly smaller than controls, we also assessed the dynamics of branch lengthening and widening in these mutants. Strikingly, our data reveal that the rates of both are significantly decreased in inducible-mesCKO lungs: bronchi from inducible-mesCKO lungs are shorter and narrower than their control counterparts at the same stage of branching (**Fig. 4I-L**). Bronchi from inducible-mesCKO lungs are also physically collapsed, lacking the clear lumenal space observed in controls (**Supplemental Fig. 4C**). These decreases in branch initiation, length, and width do not appear to result from changes in cell proliferation or death: we observe no significant differences in the percentage of phospho-histone-3-positive cells in the epithelium or mesenchyme of control and inducible-mesCKO lungs at *E*13-14 and levels of cleaved caspase-3 signal in control and inducible-mesCKO lungs at *E*13-14 are negligible (**Supplemental Fig. 4D-G**). These data demonstrate that the initiation, elongation, and widening of airway epithelial branches require expression of *Vangl* in the pulmonary mesenchyme. The pulmonary mesenchyme is therefore essential for shaping the fractal structure of the airway epithelial tree.

### Loss of mesenchymal Wnt5a does not recapitulate loss of mesenchymal Vangl1/2

Both the core PCP complex generally and Vangl specifically have been hypothesized to function downstream of non-canonical Wnt signaling, though direct evidence linking Wnt ligand binding to changes in PCP protein function is sparse. However, Wnt5a has been shown to regulate phosphorylation of Vangl2, leading to changes in cell shape and tissue elongation in the chondrocytes of the developing limb (Gao et al., 2011). Further, loss of Wnt5a and Vangl2 have similar phenotypes during late lung development, leading to speculation that a Wnt5a/Vangl signaling pathway regulates alveologenesis (Zhang et al., 2020). *Wnt5a* transcript is expressed in the mesenchymal compartment during branching morphogenesis at *E*13.5 (**Supplemental Fig. 5A**). Therefore, to determine whether Wnt5a functions upstream of mesenchymal Vangl2 in branch initiation, elongation, and widening, we generated mesenchymal knockouts of *Wnt5a*. We began by establishing a *Dermo1Cre; Wnt5a^fl/fl^* mouse line and assessed branching in control and mutant lungs at *E*12-14 (**Supplemental Fig. 5B**). Broadly, this analysis revealed that mutant lungs are misshapen and show alterations in the angles of their airways. Surprisingly, however, we observe no decrease in the number of branches in mutant lungs as compared to controls (**Supplemental Fig. 5C**). Further, *Dermo1Cre; Wnt5a^fl/fl^* embryos appear to have axis-elongation defects and exhibit broad developmental abnormalities, likely due to loss of *Wnt5a* in all *Dermo1*-expressing tissues. Thus, the *Dermo1Cre* driver cannot be used to uncouple the lung phenotypes caused by deleting *Wnt5a* from the pulmonary mesenchyme from those caused by deleting *Wnt5a* from other tissues.

To specifically uncover the role of *Wnt5a* in the lung mesenchyme, we generated a second mesenchymal *Wnt5a* knockout driven by *Tbx4-rtTA; Tet-O-Cre*. In contrast to *Dermo1Cre; Wnt5a^fl/fl^* lungs, *Tbx4-rtTA; Tet-O-Cre; Wnt5a^fl/fl^* lungs are indistinguishable from controls at *E*12 (**Supplemental Fig. 5D**). Therefore, the change in branch angle observed in *Dermo1Cre; Wnt5a^fl/fl^*lungs is likely due to loss of *Wnt5a* from other tissues, rather than due to a role for *Wnt5a* in the pulmonary mesenchyme. Strangely, *Tbx4-rtTA; Tet-O-Cre; Wnt5a^fl/fl^* knockout lungs show a slight increase in the number of branches at *E*13, but a slight decrease in the number of branches at *E*14 (**Supplemental Fig. 5E**). However, morphometric analysis reveals no significant defects in elongation or widening in the mutant lungs (**Supplemental Fig. 5F-I**). Thus, removing Wnt5a from the pulmonary mesenchyme fails to recapitulate the branching defects observed in *Vangl1/2* inducible-mesCKO lungs. While Wnt5a may regulate Vangl in this developmental context, this ligand is unlikely to be the only upstream regulator. In this regard, it is possible that the expression of another non-canonical *Wnt*, such as *Wnt11*, compensates for loss of *Wnt5a*.

### Loss of mesenchymal Vangl affects assembly of the airway smooth muscle layer

To understand how loss of mesenchymal *Vangl* leads to defects in branching morphogenesis of the adjacent epithelium, we characterized the cellular and noncellular components of the mesenchyme in control and inducible-mesCKO lungs. Immunofluorescence analysis revealed no significant differences in the expression or localization of fibronectin or collagen IV, indicating the changes observed in inducible-mesCKO lungs are unlikely to be due to alterations in the deposition of interstitial matrix or basement membrane (**Supplemental Fig. 4H-I**). Similarly, RNAscope analysis revealed no differences in mesenchymal expression of *Fgf10*, a key regulator of branching in the mouse lung (Yuan et al., 2018) (**Supplemental Fig. 4J**).

The embryonic pulmonary mesenchyme is compartmentalized into spatially distinct subpopulations of cells that express different markers and differentially affect epithelial morphogenesis; these compartments include the sub-epithelial mesenchyme, sub-mesothelial mesenchyme, and airway smooth muscle (ASM) (Goodwin et al., 2022). Immunofluorescence analysis of *E*13 inducible-mesCKO lungs revealed no changes in the distribution of sub-epithelial or sub-mesothelial mesenchyme, as assessed by the expression of Lef1 (sub-epithelial mesenchyme) and Foxp1 (sub-mesothelial mesenchyme) respectively (**Fig. 5A-D**). These data suggest that *Vangl* is not required for expression of mesenchymal markers or morphogens, nor for the patterning of mesenchymal subpopulations, consistent with the primarily structural role of this PCP component in other tissue contexts (Boutin et al., 2014, Cetera et al., 2017, Devenport and Fuchs, 2008, Gao et al., 2011).

**Figure 5.**
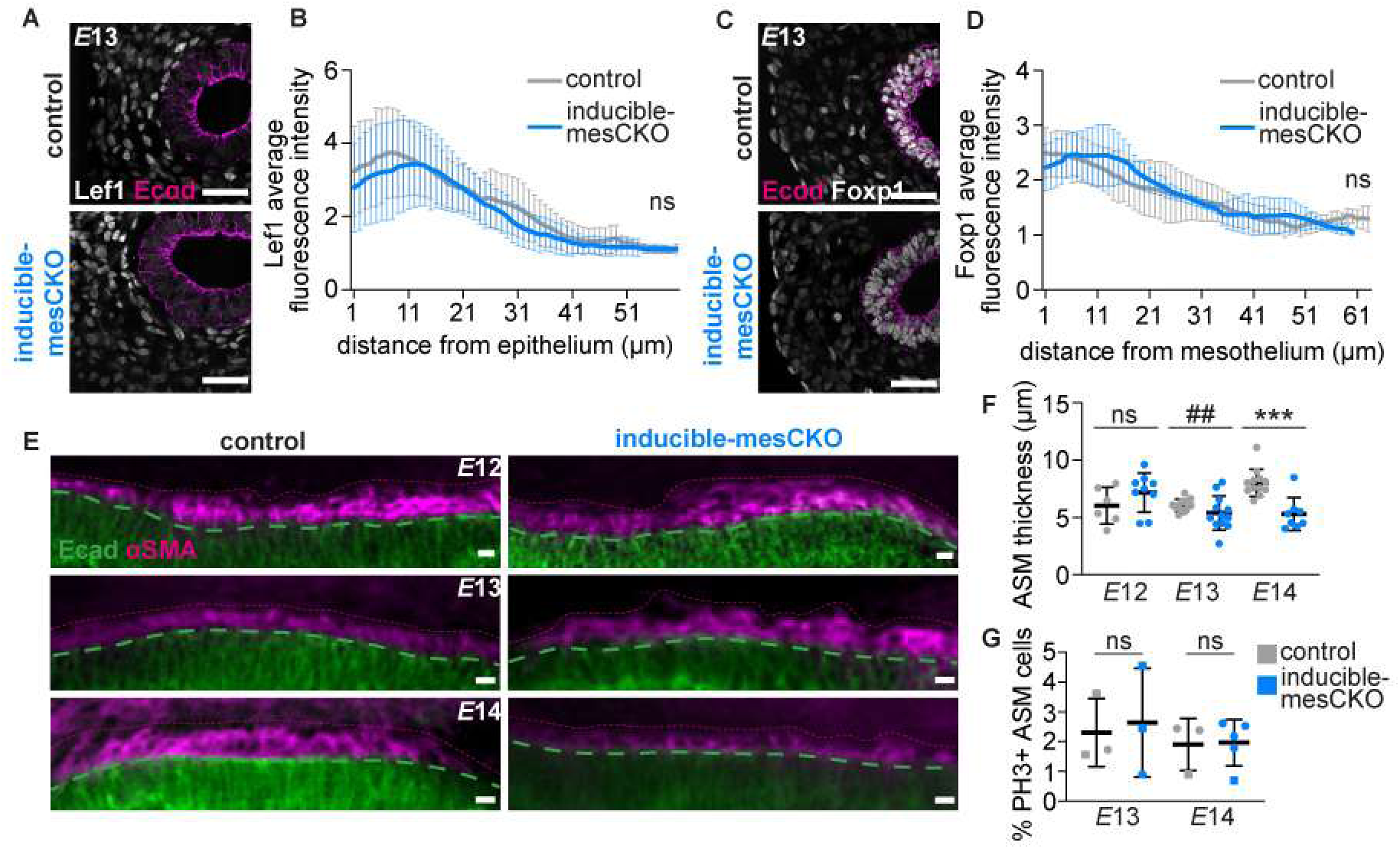
Loss of mesenchymal *Vangl* affects assembly of the airway smooth muscle layer. **A, C,** Representative images of sections (10-µm-thick) from *E*13 control and inducible-mesCKO lungs immunostained for Lef1 (sub-epithelial mesenchyme) or Foxp1 (sub-mesothelial mesenchyme). Scale bars, 50 µm. **B, D,** Quantifications of Lef1 or Foxp1 intensity profile emanating from the epithelium or mesothelium. Shown are mean ± s.d. (n = 7 control, 4 mutant). **E,** Representative image of an optical cross-section through the medial bronchi of *E*12-14 control and inducible-mesCKO lungs immunostained for Ecad and ⍺-smooth muscle actin (⍺SMA) (scale bars: 10 µm). **F,** Quantification of the thickness of the ASM layer around major bronchi at *E*12 (*n*=8 control and *n*=9 mutant bronchi, *p=*0.2105 via unpaired Student’s t-test), *E*13 (*n*=12 control and *n*=12 mutant bronchi, *p=*0.0017 via F test to compare variances), and *E*14 (*n*=12 control and *n*=9 mutant lungs, *p=*0.0002 via unpaired Student’s t-test). Shown are mean ± s.d; ## variance < 0.01; *** p < 0.001. **G,** Quantification of the percentage of phospho-histone-3^+^ (PH3^+^) ASM cells in control and inducible-mesCKO lungs at *E*13 (*n*=3 control and *n*=3 mutant lungs, *p=*0.8031 via unpaired Student’s t-test) and 14 (*n*=3 control and *n*=5 mutant lungs, *p=*0.9277 via unpaired Student’s t-test) lungs. Shown are mean ± s.d; different shapes represent different experimental replicates.

In the developing mouse lung, the sub-epithelial mesenchyme differentiates into ASM in a pattern that actively sculpts the epithelium into branches (Goodwin et al., 2019, Kim et al., 2015, Goodwin et al., 2022, Goodwin et al., 2023). To assess whether loss of Vangl leads to defects in ASM, we examined this mesenchymal layer using immunofluorescence analysis for ⍺-smooth muscle actin (⍺SMA). At *E*12, we observed no differences in the thickness of the ASM surrounding the airways between control and inducible-mesCKO lungs (**Fig. 5E-F**). At *E*13, however, we found that control ASM exists as a uniform layer approximately 5-µm thick, whereas the mutant ASM is highly variable, ranging from 2.5 to 8 µm in thickness (**Fig. 5E-F**). At *E*14, the *Vangl*-mutant smooth muscle has resolved into a layer that is significantly thinner than that of the control (**Fig. 5E-F**). This thinner layer of smooth muscle does not result from a reduction in cell proliferation (**Fig. 5G**). Taken together, these data reveal that although mesenchymal cell identity and compartmentalization do not appear to be altered in mesenchymal *Vangl*-mutants, the absence of this protein results in defects in the organization of cells within the ASM tissue layer.

### Loss of mesenchymal Vangl alters the morphology of airway smooth muscle cells

In most other developmental and cellular contexts, Vangl predominately regulates cellular behaviors, such as cell motility, shape, and polarization, rather than transcription. Given the differences in thickness of the ASM layer, we hypothesized that mesenchymal Vangl may affect epithelial morphogenesis by regulating the morphology of ASM cells. We generated a *Tbx4-rtTA; Tet-O-Cre; Vangl1^fl/fl^; Vangl2^fl/fl^; Confetti* mouse line, which enabled us to analyze the shapes of sparsely labeled *Vangl*-mutant smooth muscle cells. In control (*Tbx4-rtTA; Tet-O-Cre; Confetti*) lungs, ⍺SMA^+^ cells are highly elongated and wrap circumferentially around the airways (**Fig. 6A**). However, in inducible-mesCKO lungs, ⍺SMA^+^ cells appear rounder and less elongated (**Fig. 6B**). Cell-shape analysis revealed that *Vangl*-mutant ASM cells have a significantly reduced aspect ratio and increased circularity (**Fig. 6C-D**). Further, *Vangl*-mutant ASM cells exhibit a reduction in circumferential polarization around the airway epithelium (**Fig. E-G**). Therefore, *Vangl* is required for the normal elongation and circumferential wrapping of ASM cells in the developing lung. These changes in smooth muscle morphology coincide with delays in branch initiation, lengthening and widening in the airways of the *Vangl*-mutant animals.

**Figure 6.**
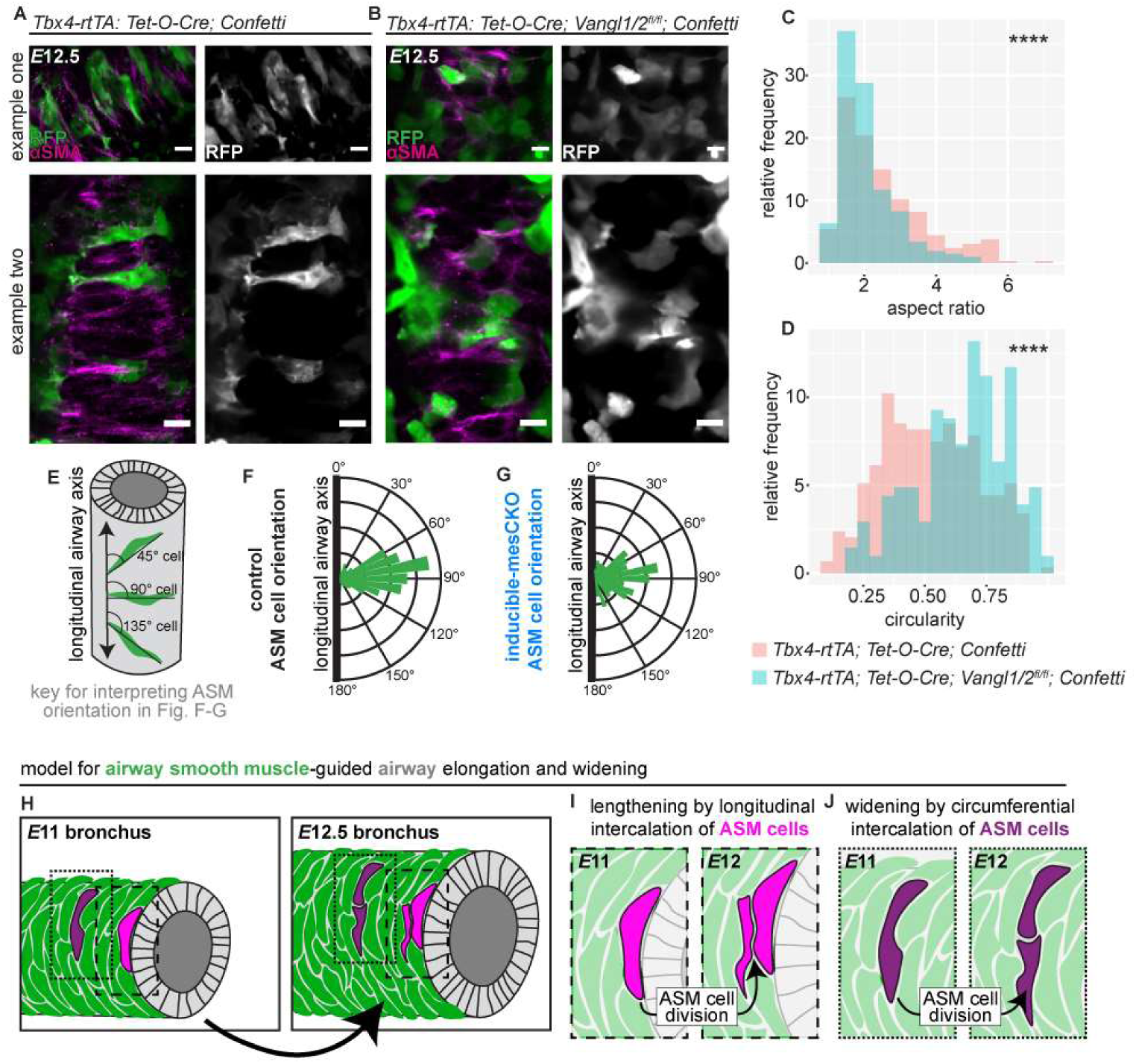
Loss of mesenchymal *Vangl1/2* alters the morphology of ASM cells. **A,** Examples of ASM cells [immunostained for ⍺SMA (magenta) with signal from RFP fluorescence (green)] in *Tbx4-rtTA; Tet-O-Cre; Confetti* lungs (scale bars: 10 µm). **B,** Examples of ASM cells [immunostained for ⍺SMA (magenta) with signal from RFP fluorescence (green)] in *Tbx4-rtTA; Tet-O-Cre; Vangl1/2^fl/fl^; Confetti* lungs (scale bars: 10 µm). **C-D,** Histograms of the aspect ratio (p < 0.0001, Mann-Whitney test) and circularity (p < 0.0001, Mann-Whitney test) of ASM cells from *E*12.5 *Tbx4-rtTA; Tet-O-Cre; Confetti* (n = 294 cells from 3 lungs) and *Tbx4-rtTA; Tet-O-Cre; Vangl1^fl/fl^; Vangl2^fl/fl^; Confetti* lungs (n = 205 cells from 4 lungs). **** p < 0.0001. **E,** Cartoon illustrating how ASM cell orientations were measured relative to the long axis of the airway. **F,** Rose plot of ASM cell orientations from control *E*12.5 lungs (n =296 cells from 3 lungs). **G,** Rose plot of ASM cell orientations from inducible-mesCKO *E*12.5 lungs (n =224 cells from 4 lungs). **H,** Cartoon schematic of proposed model for ASM-regulated widening and lengthening of airways. **I,** Schematic of how intercalation of newly-divided ASM cells along the circumferential axis of the airway lead to widening. **J,** Schematic of how intercalation of newly divided ASM cells along the longitudinal axis of the airway lead to lengthening.

### Vangl2 is broadly expressed throughout the mesenchyme in organogenesis

Our data show that Vangl1/2 function outside of the core PCP complex in an intricate, nonplanar, 3D tissue to facilitate mesenchymal organization and epithelial morphogenesis. We hypothesize that Vangl exerts control over the cytoskeleton to correctly structure tissue layers within the mesenchyme. We were curious whether this function for Vangl in the mesenchyme might be conserved in other organs.

To assess whether there may be a broader role for *Celsr1*-independent *Vangl2* in organogenesis, we took advantage of an endogenously tagged *Vangl2-tdTomato* mouse line to examine the expression pattern of Vangl2 across various organs (Basta et al., 2021). Our analysis revealed that Vangl2 is localized to the membrane in mesenchymal cells of the developing kidney and intestine (**Fig. 7A**). Consistent with our data for the lung mesenchyme, scRNA-seq analysis shows that *Vangl2* is expressed ubiquitously while *Celsr1* is excluded from the mesenchyme in these organs (**Fig. 7B-C, Supplemental Fig. 7A-B**) (Naganuma et al., 2021, Dong et al., 2018). Thus, Vangl2 is expressed in the mesenchyme in the absence of the key core PCP component Celsr1 in several embryonic mouse organs.

**Figure 7.**
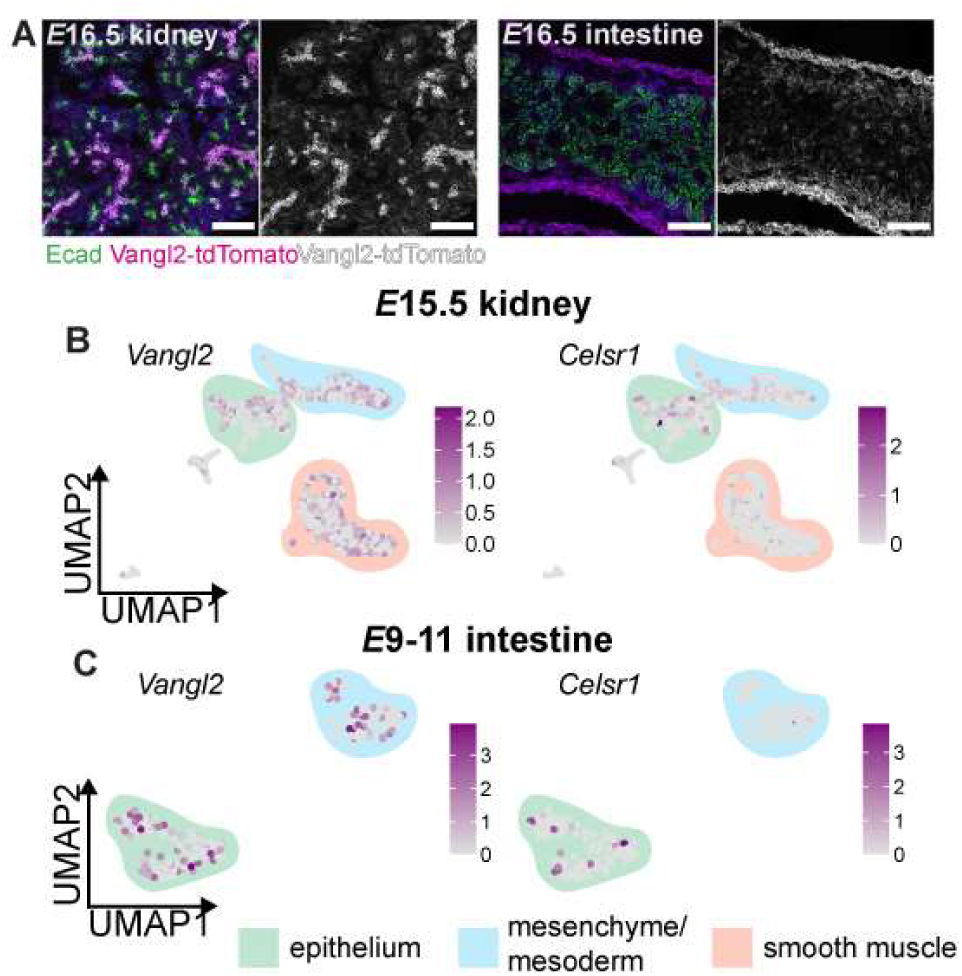
Vangl2 is widely expressed in embryonic mesenchymal tissues. **A,** Representative images of sections (10-µm-thick) from *E*16.5 *Vangl2-tdTomato* kidney and intestine immunostained for Ecad (green) or tdTomato (magenta) with nuclei counterstained with Hoechst (blue). Scale bars, 50 µm. **B-C,** scRNA-seq of developing kidney or intestine revealing expression of *Vangl2* or *Celsr1* in tissue compartments (Naganuma et al., 2021, Dong et al., 2018).

## Discussion

While decades of research have sought to understand how new branches initiate during morphogenesis of the mouse lung, we still lack a clear understanding of how the embryonic airways increase in size to form a mature, fractal-shaped organ. Our data reveal that neither *Vangl1/2* nor *Celsr1*, core members of the PCP complex, are necessary in the epithelium for branch initiation, elongation, or widening during the pseudoglandular stage of lung development. Previous work concluded that the number of branches is reduced in *Vangl2^Lp/Lp^*and *Celsr1^Crsh/Crsh^* lungs (Yates et al., 2010). However, we observe no reduction in branching in *Celsr1^Crsh/Crsh^* lungs *in vivo* or in cultured lung explants. Further, the reduction in branching observed in *Vangl2^Lp/Lp^* lungs can be attributed entirely to loss of *Vangl* from the mesenchymal compartment, whereas epithelial *Vangl* is dispensable for branch initiation, elongation, and widening.

Our work contradicts conclusions from a recent study, which reported a slight alteration in the locations and angles of epithelial branches when *Vangl1/2* are globally removed early in mouse embryonic development (using *Sox2Cre; Vangl1^gt/gt^; Vangl2^fl/fl^* animals) (Zhang et al., 2022). We observe no obvious changes in airway location or angle in either *Vangl2^Lp/Lp^* or *Vangl1/2* inducible-mesCKO lungs. Indeed, we observe a distal tilting of the cranial, medial, and L1 bronchi in these mutant lungs by *E*13, comparable to controls. In the same study, the authors also attempted to assess tissue-specific functions of *Vangl1/2* by using global epithelial (*Sox9Cre*) and mesenchymal (*Dermo1Cre*) drivers. In agreement with our experiments, the authors observed no phenotype in either of these lines. However, while the authors interpreted this absence of phenotype to mean that *Vangl1/2* are required in both the epithelium and the mesenchyme, we find that fully removing *Vangl1/2* only from the mesenchyme results in defects in branching morphogenesis.

Our study reveals the exciting result that *Vangl1/2* expressed specifically in mesenchymal cells plays a role outside of the core PCP complex to promote airway initiation, elongation, and widening, processes that are essential for generating the fractal architecture of the mature epithelial tree. Lungs lacking mesenchymal *Vangl* show clear defects in airway elongation and widening by *E*14 and exhibit a severe reduction in branch initiation at *E*14. This represents a new function for Vangl outside of its canonical role in the PCP complex, as the core PCP components Celsr1-3 are absent from the pulmonary mesenchyme. Wnt5a has been hypothesized to function upstream of both the core PCP complex generally and Vangl2 specifically in many tissues. Our data show that specific loss of *Wnt5a* from the pulmonary mesenchyme does not affect widening or elongation, though we do observe a slight decrease in branch initiation by *E*14. In fact, mesenchymal *Wnt5a* knockout lungs appear to be phenotypically normal at *E*12, in contrast to recent work concluding that loss of mesenchymal *Wnt5a* leads to an alteration in the position and angle of branches (Zhang et al., 2022). Though broad deletion of splanchnic-mesodermal *Wnt5a* driven by *Dermo1Cre* does cause gross defects in the morphology of the lung, we do not observe the same phenotype when *Wnt5a* is specifically removed from the pulmonary mesenchyme using *Tbx4-rtTA; Tet-O-Cre*. This difference in phenotype can likely be attributed to the expression pattern of these Cre lines: *Dermo1Cre* is expressed in tissues outside of the lung, while the expression of *Tbx4-rtTA; Tet-O-Cre* is restricted to the pulmonary mesenchyme. Given that branching patterns are altered when the lung is constrained (Nelson et al., 2017, Gilbert et al., 2021), we speculate that loss of *Wnt5a* in *Dermo1Cre*-expressing non-pulmonary tissues leads to changes in the shape of the chest cavity, thereby stunting morphogenesis.

Our morphometric analysis revealed that the defects in airway lengthening observed in *Vangl1/2* inducible-mesCKO lungs occurred concomitantly with defects in airway widening. Specifically, airways from inducible-mesCKO lungs are both shorter and narrower than their wildtype counterparts. This phenotype was initially surprising, as defects in tissue elongation are often coupled with abnormal widening along the orthogonal axis; for example, embryos that fail to undergo convergent extension are short and wide (Goto and Keller, 2002, Paudyal et al., 2010, Sutherland et al., 2020). However, this paradoxical phenotype has been observed previously in other mutations that affect smooth muscle. For example, loss of the potassium channel KCNJ13 leads to disorganized smooth muscle alignment around the trachea and both shorter and narrower tracheas in mice (Yin et al., 2018). Similarly, in *Ror2-* and *Wnt5a*-mutant lungs, tracheal smooth muscle cells exhibit defects in polarization and radial intercalation, resulting in short and narrow tracheas (Kishimoto et al., 2018).

At the cellular level, we find that loss of *Vangl1/2* leads to alterations in the morphology of ASM cells. In control lungs, ASM cells elongate and circumferentially wrap airways. When *Vangl* is absent, ASM cells do not elongate and instead exhibit a cobblestone-like morphology. We hypothesize that normal elongation and wrapping of smooth muscle cells is required for airway elongation and widening. The ASM layer is a stiffer tissue than both the airway epithelium and the surrounding undifferentiated mesenchyme (Goodwin et al., 2020, Goodwin et al., 2023). We propose a model wherein the ASM layer functions as a brace, preventing inward collapse of the airway through the adhesion of the epithelium to their shared basement membrane. We hypothesize that continual axial and circumferential intercalation of newly-divided ASM cells along established airways ratchets the epithelial tube both longer and wider (**Fig. 6H-J**). We have termed this proposed mechanism ‘dilational extension,’ in contrast to the convergent-extension mechanism that lengthens epithelia in other branched organs such as the kidney (Kunimoto et al., 2017, Lienkamp et al., 2012). A similar set of mesenchymal rearrangements is thought to play a role in the developing mouse trachea: the polarization and intercalation of tracheal smooth muscle cells leads to both tracheal elongation and widening (Kishimoto et al., 2018). We hypothesize that Vangl regulates the polarization of ASM cells, and thus their ability to intercalate and wrap properly around the airways. Without the requisite continual intercalation of new ASM cells, we posit that the ASM tissue layer is organized improperly and cannot exert the forces required to promote elongation and widening of the airways.

Our data suggest that the PCP complex plays a distinct role in the lung as compared to other branched organs, such as the kidney and mammary gland. PCP is not required in the lung epithelium for the initiation or extension of branches, unlike the significant defects observed in epithelial-specific PCP-mutant kidneys and mammary glands (Lienkamp et al., 2012, Smith et al., 2019, Kunimoto et al., 2017). Rather, the key role for the PCP complex in the airway epithelium appears to be generation and maintenance of cilia polarity (Vladar et al., 2012, Kunimoto et al., 2022, Vladar et al., 2016). Our data thus indicate that the PCP complex is not universally required in the epithelium for the polarized behaviors that contribute to branching morphogenesis.

It is likely that Vangl2 has PCP-independent functions in organs beyond the lung. We find that Vangl2 is expressed in the absence of Celsr1 in both the kidney and intestinal mesenchyme, suggesting a role outside of the core PCP complex in those tissues. A related polarity pathway, the Fat-Dchs pathway, has been shown to interact genetically with Vangl2 to regulate the migration and clustering of mesenchymal cells in the developing intestine (Rao-Bhatia et al., 2020). Collectively, these data suggest a role for polarity signaling in diverse, 3D mesenchymal tissues. We propose that, in the context of vertebrate development, Vangl has evolved functions distinct from the core PCP complex to regulate mesenchymal cell dynamics during organogenesis. Future work will explicitly uncover the pathway via which mesenchymal Vangl2 functions to promote epithelial remodeling to form the fractal tree of airways.

## Materials and Methods

### Mouse lines and breeding

All procedures involving animals were approved by Princeton University’s Institutional Animal Care and Use Committee (IACUC). Mice were housed in an AAALAC-accredited facility in accordance with the NIH Guide for the Care and Use of Laboratory Animals. This study was compliant with all relevant ethical regulations regarding animal research. CD1 mouse embryos were used for scRNA-seq (Goodwin et al., 2022) and immunostaining experiments to determine expression and localization of PCP components. *Vangl2^Lp/Lp^*embryos (Kibar et al., 2001) and *Celsr1^Crsh/Crsh^* embryos (Curtin et al., 2003) were used to determine how the loss of PCP function affects airway epithelial branching. *ShhCreGFP; Vangl1^fl/fl^; Vangl2 ^fl/fl^; Rosa26mTmG/Rosa26mTmG* embryos were used to conditionally delete *Vangl1/2* from the lung epithelium (Harfe et al., 2004, Harris et al., 2006, Copley et al., 2013). *Dermo1Cre; Vangl1 ^fl/fl^; Vangl2 ^fl/fl^; Rosa26mTmG/Rosa26mTmG* embryos were used to conditionally delete *Vangl1/2* from the pulmonary mesenchyme (Yu et al., 2003, Yin et al., 2008, Sosic et al., 2003). *Tbx4-rtTA; Tet-O-Cre; Vangl1 ^fl/fl^; Vangl2 ^fl/fl^; Rosa26mTmG/Rosa26mTmG* embryos were used to conditionally delete *Vangl1/2* from the pulmonary mesenchyme in an inducible manner (Zhang et al., 2013). To induce deletion of *Vangl1/2*, doxycycline-medicated water (0.5 mg/mL) was administered to pregnant dams at *E*8. *Dermo1Cre; Wnt5a^fl/fl^*embryos were used to conditionally delete *Wnt5a* from the pulmonary mesenchyme (Ryu et al., 2013). *Tbx4-rtTA; Tet-O-Cre; Wnt5a ^fl/fl^* embryos were used to conditionally delete *Wnt5a* from the pulmonary mesenchyme in an inducible manner; doxycycline was administered in drinking water at *E*8 as described above. *Tbx4-rtTA; Tet-O-Cre; Vangl1 ^fl/fl^; Vangl2 ^fl/fl^; R26R-Confetti* (Livet et al., 2007) embryos were used for cell-shape analysis. Genotyping primers are listed in **Supplementary Material**.

### Immunofluorescence analysis

Lungs were dissected from embryos in PBS and fixed in 4% paraformaldehyde (PFA). Wholemount *E*11.5-13.5 lungs were fixed for 30 min at 4℃, while wholemount E14.5 lungs were fixed for 1 h at 4℃. Lungs were then washed in PBS and blocked overnight in blocking buffer composed of 4% normal donkey serum, 1% bovine serum albumin (BSA), and 1% fish gelatin in PBT2 (PBS with 0.2% Triton X-100). Lungs were incubated with primary antibody in blocking buffer overnight (2 nights for *E*14.5 lungs). Lungs were washed with PBS, incubated with secondary antibody overnight (2 nights for *E*14.5 lungs), and then washed in PBS, dehydrated through a methanol series, and cleared for imaging in 1:1 benzyl alcohol:benzyl benzoate (BABB). After fixation, lungs for sectioning were washed in PBS, and taken through a sucrose gradient before embedding in OCT. 10-µm-thick or 200-µm-thick frozen sections were obtained from samples using a Leica CM3050S cryostat. 10-µm-thick sections were washed in PBT3 (PBS with 0.3% Triton X-100), washed in PBS, incubated in blocking buffer for 1 h at room temperature, and then incubated in blocking buffer with primary antibody overnight. Slides were then washed in PBS and incubated with secondary antibody for 3 h at RT, washed, and mounted in Prolong Gold. 200-µm-thick floating sections were washed in PBT3, washed in PBS, incubated in blocking buffer overnight at 4℃, and then incubated in blocking buffer with primary antibody overnight. Sections were then washed in PBS and incubated with secondary antibody overnight at 4℃. Sections were then washed in PBS and mounted in Prolong Gold. All antibodies used for staining are described in **Supplementary Material**.

### Single-cell RNA-sequencing (scRNA-seq) analysis

Data for scRNA-seq analysis were generated in a previous study (Goodwin et al., 2022). Briefly, the left lobes of lungs were dissected in cold PBS from CD1 mouse embryos collected at *E*11.5, then placed in dispase and mechanically dissociated using tungsten needles. After 10 min in dispase at room temperature, DMEM without HEPES and supplemented with 5% fetal bovine serum (FBS, Atlanta Biologicals) was added, and cell suspensions were passed through a filter with 40-µm-diameter pores. The resultant cell suspensions were then processed by the Princeton Genomics Core Facility, using the Chromium Single Cell 3’ Library and Gel Bead Kit v2, and then loaded onto the Chromium Controller (10X Genomics). Illumina sequencing libraries were prepared from the amplified cDNA from each sample group using the Nextra DNA library prep kit (Illumina) and then sequenced on Illumina HiSeq 2500 Rapid flowcells (Illumina) as paired-end 26 + 50 nucleotide reads. Reads were processed using the Illumina sequencer control software to retain only pass-filtered reads for downstream analysis. The 10× CellRanger software version 2.0.1 was used to generate gene-barcode matrices using the *Mus musculus* reference genome mm10-1.2.0.

Data were then processed using the Seurat package in R following standard procedures (Butler et al., 2018). The data were normalized, integrated based on 2000 variable genes, then scaled and analyzed to find neighbors and clusters. Clusters were identified based on known cell-type markers, and gene expression in each cell type was visualized using violin plots.

We analyzed two additional published scRNA-seq datasets generated from embryonic mouse tissues: *E*15.5 kidney (GSE149134) (Naganuma et al., 2021) and *E*9.5-11.5 intestine (GSE87038) (Dong et al., 2018). All analyses were carried out using the Seurat package (Butler et al., 2018). The intestinal scRNA-seq data consist of four datasets from embryos at different stages of development (two at *E*9.5, one at *E*10.5, and one at *E*11.5), which we merged prior to analysis. The data were first filtered to exclude cells with fewer than 500 genes, more than 30,000 unique molecular identifiers (possible multiplets), or greater than 10% mitochondrial DNA (dying cells). Following the Seurat pipeline, we then normalized the data, identified variable features, scaled gene expression for each cell, ran principal components analysis, and identified neighbors and clusters. We then generated uniform manifold approximation and projection (UMAP) plots and extracted cluster markers to identify cell types. Clusters representing distinct cell types were identified by color-coding the UMAP based on known markers of epithelial and mesenchymal cells in the kidney (*Cdh1* and *Cdh2*) and the intestine (*Cdh1* and *Foxf1*). We then examined the expression patterns of *Celsr1* and *Vangl2* by color-coding the UMAP and comparing cell-level expression of these two genes to that of the cell-type-specific markers listed above.

### Organ explant culture

*E*12.5 lungs were dissected in sterile PBS and cultured at an air-liquid interface on porous membranes (nucleopore polycarbonate track-etch membrane, 8-μm pore size, 25-mm diameter; Whatman) in DMEM/F12 medium (without HEPES) supplemented with 5% FBS and antibiotics (50 units/ml of penicillin and streptomycin). Explants were fixed for 30 min in 4% PFA at room temperature and stained and imaged as described above.

### Fluorescence in situ hybridization analysis

Lungs were isolated at *E*12.5-13.5 and immediately fixed in 4% PFA in DEPC-treated PBS for 24 h at 4°C. Lungs were taken through a sucrose gradient (20% sucrose in DEPC-treated PBS for 1 h at 4°C, then 30% sucrose in DEPC-treated PBS overnight at 4°C), embedded in OCT (Tissue Tek), and frozen on dry ice. 10-µm-thick frozen sections were obtained from each sample using a Leica CM3050S cryostat. Fluorescence *in situ* hybridization was performed using the standard RNAscope Multiplex Fluorescent V2 Assay (ACD) protocol for fixed-frozen samples. The probes were for *Mus musculus Fgf10* (channel 2, 446371) and *Wnt5a* (channel 2, 316791) and the fluorophores were Opal Polaris 520 and Opal 620 (Akoya Biosciences FP1487001KT and FP1495001KT). Sections were imaged on a Nikon A1RSi confocal microscope with a 40× objective.

## Acknowledgements

This work was supported in part by grants from the National Institutes of Health (HL110335, HL118532, HL120142, HL164861, HD099030, HD111539, AR066070, AR068320), the Genetics and Molecular Biology Training Grant of the Molecular Biology Department at Princeton University (T32 GM007388), and a Faculty Scholars Award from the Howard Hughes Medical Institute. S.V.P was supported in part by a Ruth L. Kirschstein (F31) Fellowship. K.G. was supported in part by a postgraduate scholarship-doctoral (PGS-D) from the Natural Sciences and Engineering Research Council of Canada, the Dr. Margaret McWilliams Predoctoral Fellowship from the Canadian Federation of University Women, a predoctoral fellowship from the American Heart Association, and the Princeton University Procter Fellowship. We thank Dr. Wei Shi (Keck School of Medicine of USC) for generously providing us with the *Tbx4-rtTA; Tet-O-Cre* mice. We would also like to thank Sha Wang and the Microscopy Core Facility at Princeton University, a Nikon Center for Excellence for their assistance with imaging. We also thank Katie Little for her huge assistance in genotyping and maintaining the mouse colonies used in this work.

---

**Supplemental Figure 1.**
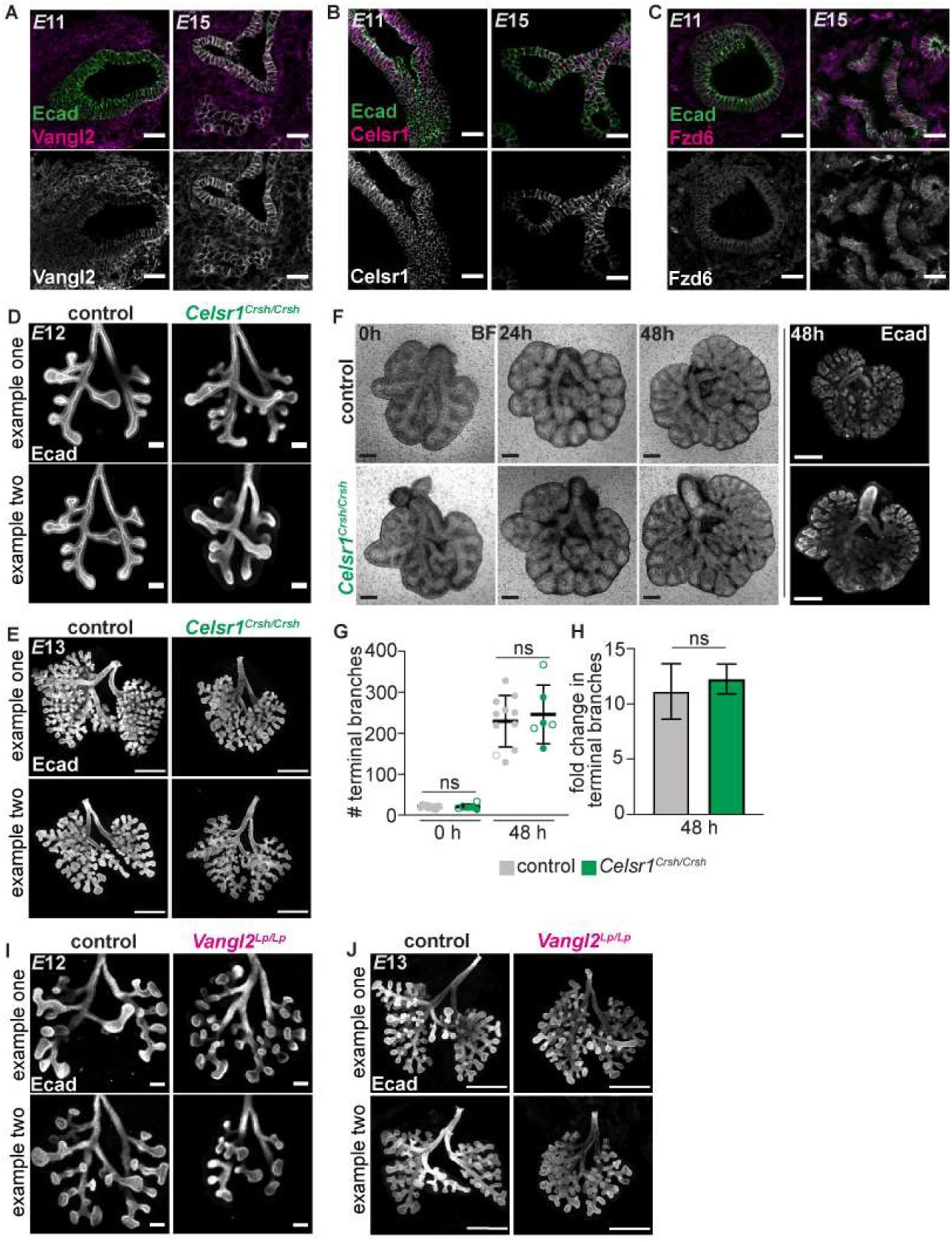
Celsr1 is dispensable for branching morphogenesis. **A-C,** Sections (10-µm-thick) from *E*11 and *E*15 lungs immunostained for Vangl2, Celsr1, or Fzd6 (magenta) and Ecad (green). Vangl2 is at the membrane in epithelial and mesenchymal cells while Celsr1 is restricted to epithelial cells; scale bars, 25 µm. **D-E,** Z-projections of confocal slices acquired from cleared, whole-mount lungs from *E*12 (scale bars, 100 µm) and *E*13 (scale bars, 500 µm) control and *Celsr1^Crsh/Crsh^* embryos immunostained for Ecad. **F,** Representative time-lapse images of lungs dissected from control and *Celsr1^Crsh/Crsh^* embryos at *E*12 and cultured for 48 h; scale bars, 250 µm. **G,** Number of terminal branches in lungs dissected from control and *Celsr1^Crsh/Crsh^* embryos after 0 h and 48 h of culture (*n*=11 control and *n*=6 mutants, *p*=0.9300 at 0 h, *p*=0.6258 at 48 h via unpaired Student’s t-test). **H,** Fold change in number of terminal branches after 48 h of culture for lungs dissected from control and *Celsr1^Crsh/Crsh^* embryos (*n*=11 control and *n*=6 mutants, *p*=0.3270 via unpaired Student’s t-test). **I-J,** Z-projections of confocal slices acquired from cleared, whole-mount lungs from *E*12 (scale bars, 100 µm) and *E*13 (scale bars, 500 µm) control and *Vangl2^Lp/Lp^* embryos immunostained for Ecad.

**Supplemental Figure 2.**
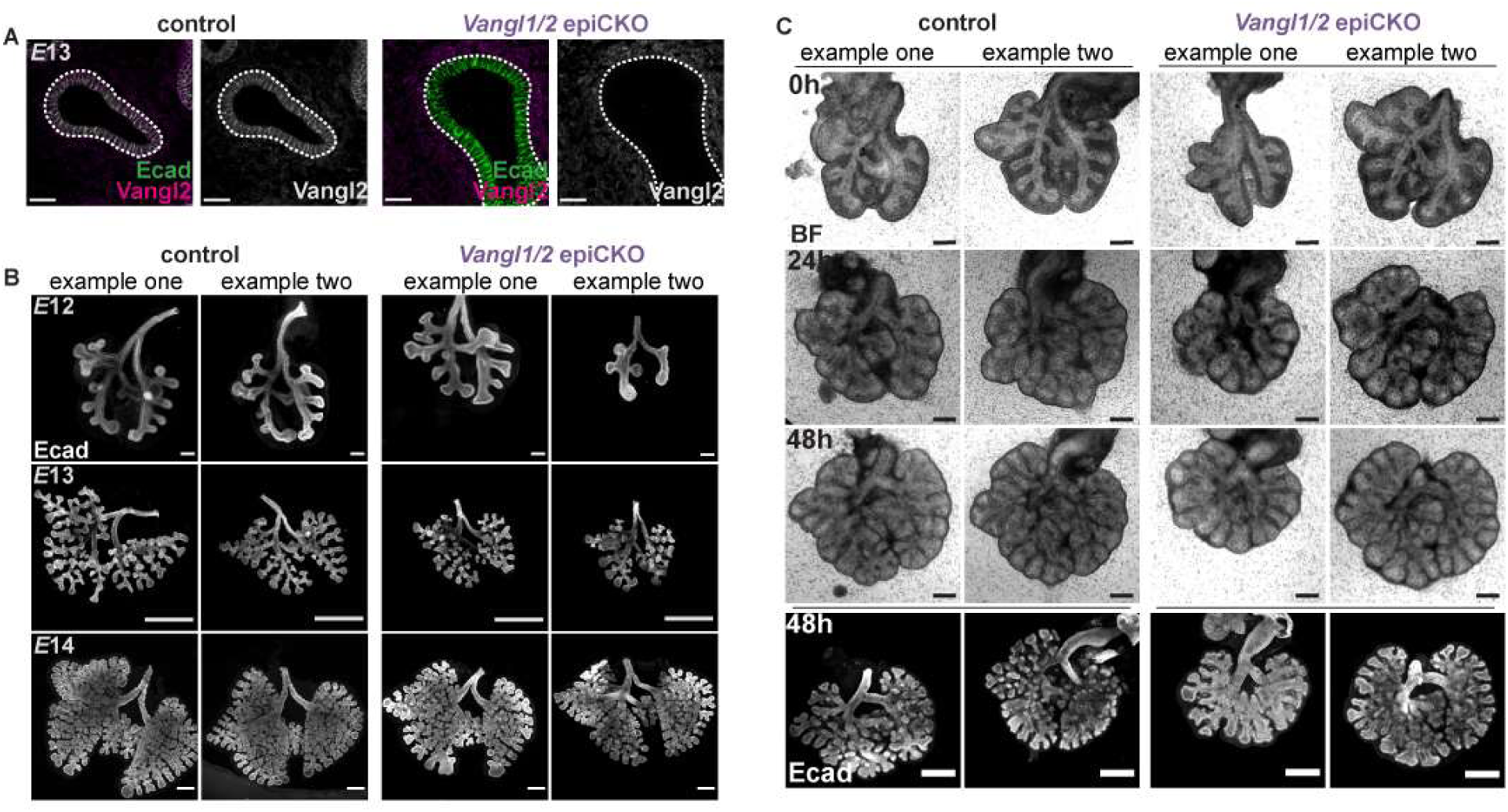
Loss of detectable Vangl2 protein in the lung epithelium using the *ShhCre* driver does not impair branch initiation. **A,** Sections (10-µm-thick) from *E*13 control and epiCKO lungs immunostained for Vangl2 (magenta) and Ecad (green), showing loss of Vangl2 from the epithelial compartment. Dashed lines outline the epithelium; scale bars, 25 µm. **B,** Z-projections of confocal optical slices acquired from cleared, whole-mount *E*12 (scale bars, 100 µm), *E*13 (scale bars, 500 µm), and *E*14 (scale bars, 250 µm) lungs from control and epiCKO embryos. **C**, Representative time-lapse images of control and epiCKO lungs dissected at *E*12 and cultured for 48 h; scale bars, 250 µm.

**Supplemental Figure 3.**
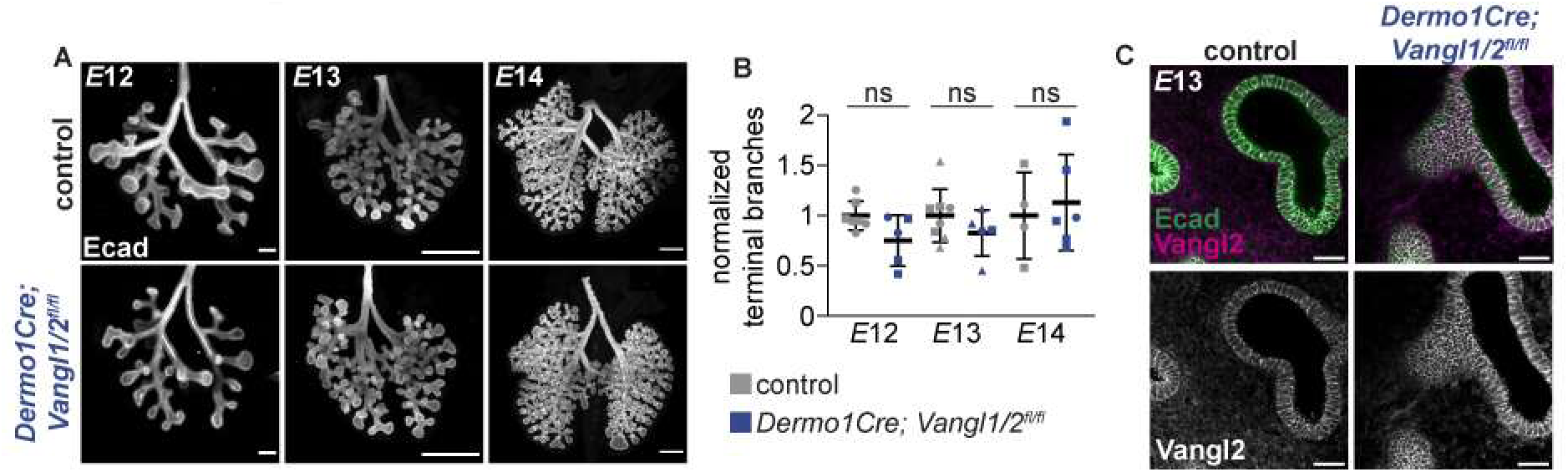
*Dermo1Cre* does not remove Vangl2 protein from the pulmonary mesenchyme. **A,** Z-projection of confocal optical slices acquired from cleared, whole-mount *E*12 (scale bars, 100 µm), *E*13 (scale bars, 250 µm), and *E*14 (scale bars, 250 µm) lungs from control and *Dermo1Cre; Vangl1/2^fl/fl^; mTmG* embryos. **B,** Normalized number of terminal branches in control and *Dermo1Cre; Vangl1/2^fl/fl^; mTmG* lungs at *E*12 (*n*=7 control and *n*=5 mutant lungs, *p*=0.0553 via unpaired Student’s t-test), *E*13 (*n*=8 control and *n*=5 mutant lungs, *p*=0.2569 via unpaired Student’s t-test), and E14 (*n*=4 control and *n*=6 mutant lungs, *p*=0.6751 via unpaired Student’s t-test). Different shapes represent different experimental replicates. **C,** Sections (10-µm-thick) from *E*13 control and *Dermo1Cre; Vangl1/2^fl/fl^; mTmG* lungs immunostained for Vangl2 (magenta) and Ecad (green), showing presence of Vangl2 in the mesenchymal compartment. Scale bars, 25 µm.

**Supplemental Figure 4.**
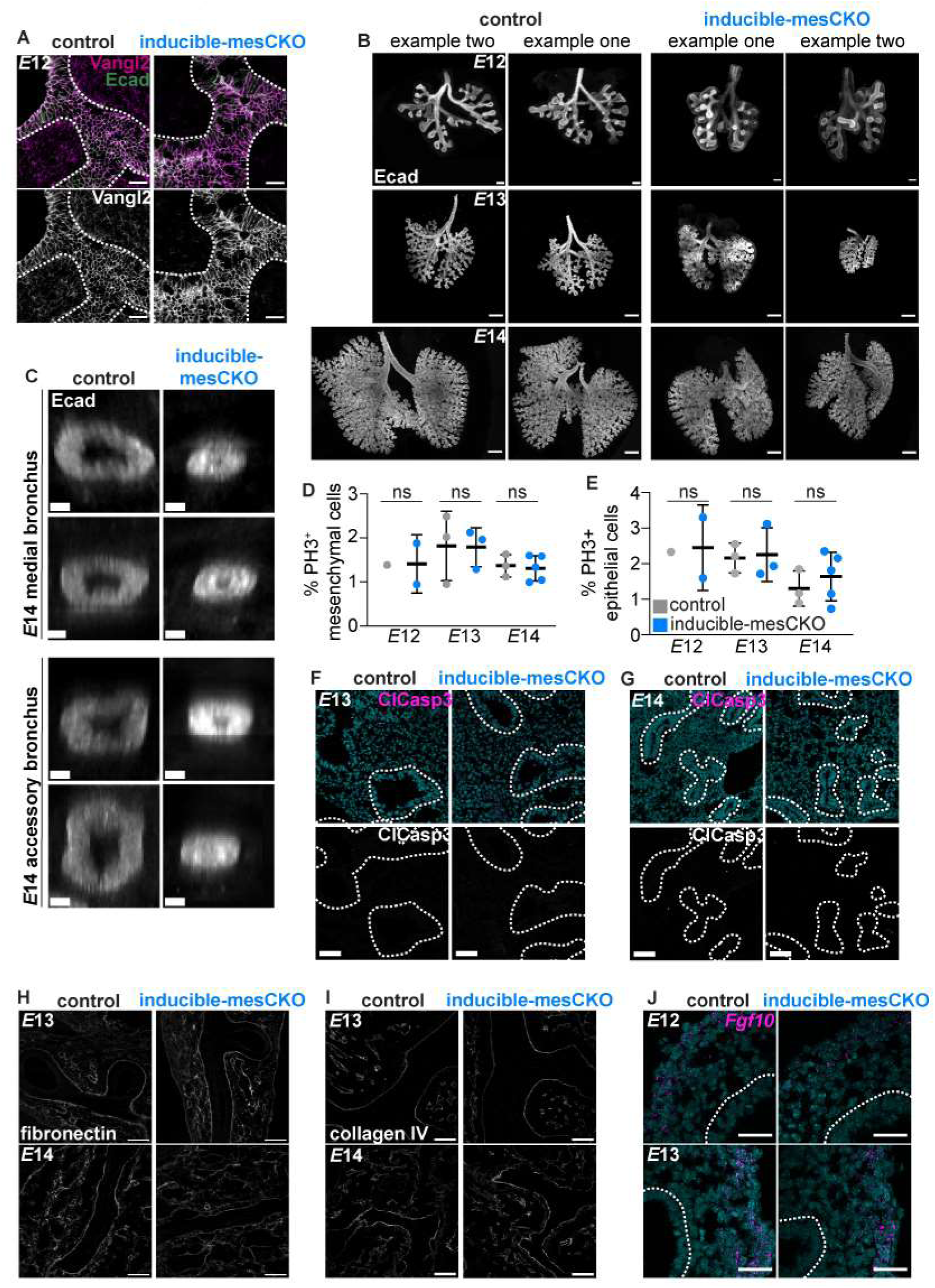
Characterization lungs from *Tbx4-rtTA; Tet-O-Cre; Vangl1^fl/fl^; Vangl2^fl/fl^*embryos. **A,** Sections (10-µm-thick) from *E*12 control and inducible-mesCKO lungs immunostained for Vangl2 (magenta) and Ecad (green), showing loss of Vangl2 from the mesenchymal compartment. Dashed lines outline the epithelium; scale bars, 25 µm. **B,** Z-projection of confocal optical slices acquired from cleared, whole-mount *E*12 (scale bars, 100 µm), *E*13 (scale bars, 250 µm), and *E*14 (scale bars, 250 µm) lungs from control and inducible-mesCKO embryos. **C,** Representative optical cross-sections of bronchi in lungs from E14.5 control and inducible-mesCKO embryos to show collapse of airways in mutants. **D-E,** Quantification of phospho-histone-3 positive (PH3^+^) mesenchymal (Ecad^-^) or epithelial (Ecad^+^) cells in control and inducible-mesCKO lungs at *E*12, *E*13 (*n*=3 control and *n*=3 mutant lungs, *p*=0.8605 for epithelium and *p*=0.9622 for mesenchyme via unpaired Student’s t-test), and *E*14 (*n*=3 control and *n*=5 mutant lungs, *p*=0.4864 for epithelium and *p*=0.7820 for mesenchyme via unpaired Student’s t-test); each dot represents one lung (four fields of view, 307 µm^2^ each) quantified per lung, dot represents the average number of proliferating cells from all images). **F-G,** Representative images from sections (10-µm-thick) from *E*13-14 control and inducible-mesCKO lungs immunostained for cleaved-caspase-3, with nuclei counterstained with Hoechst (blue). Dotted lines outline epithelium. No cleaved-caspase-3 positive cells were observed in most control or mutant lungs. Scale bars, 50 µm. **H-I,** Sections (10-µm-thick) from *E*13-14 control and inducible-mesCKO lungs immunostained for fibronectin or collagen IV (scale bars, 50 µm). **J,** RNAscope analysis of *Fgf10* transcript in *E*12-13 control and inducible-mesCKO lungs. Dotted lines outline epithelium (scale bars, 50 µm).

**Supplemental Figure 5.**
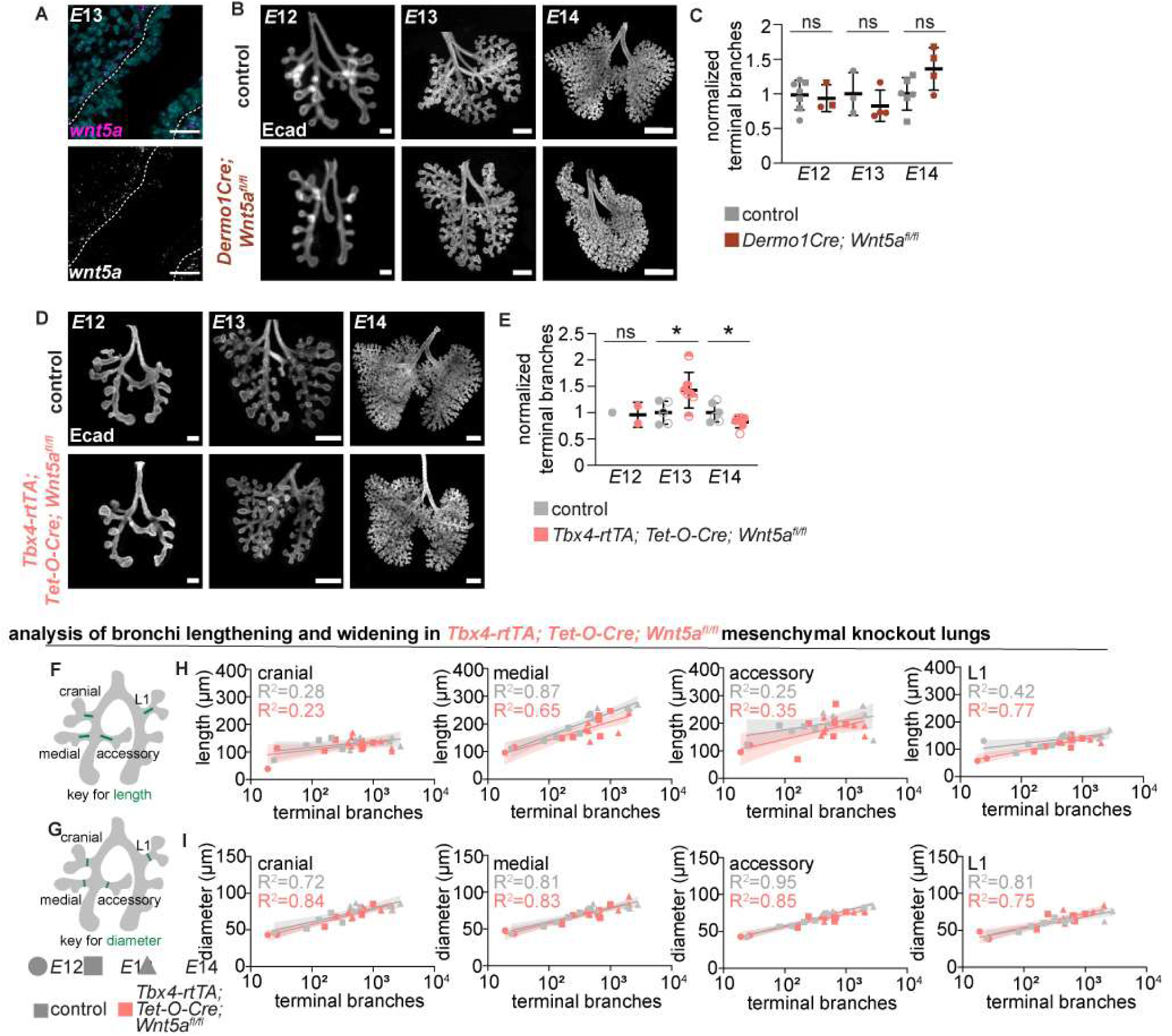
Loss of mesenchymal *Wnt5a* does not recapitulate loss of mesenchymal *Vangl1/2*. **A,** Sections (10-µm-thick) from *E*13 control lungs showing RNAscope analysis for *Wnt5a* transcript. Dotted lines outline epithelium (scale bars, 25 µm). **B,** Z-projection of confocal optical slices acquired from cleared, whole-mount *E*12 (scale bars, 100 µm), *E*13 (scale bars, 250 µm), and *E*14 (scale bars, 500 µm) lungs from control and *Dermo1Cre; Wnt5a^fl/fl^* embryos. **C,** Normalized number of terminal branches in control and *Dermo1Cre; Wnt5a^fl/fl^* lungs at *E*12 (*n*=7 control and *n*=3 mutant lungs, *p*=0.7620 via unpaired Student’s t-test), *E*13 (*n*=3 control and *n*=4 mutant lungs, *p*=0.4274 via unpaired Student’s t-test), and *E*14 (*n*=6 control and *n*=4 mutant lungs, *p*=0.0676 via unpaired Student’s t-test). **D,** Z-projection of confocal optical slices acquired from cleared, whole-mount *E*12 (scale bars, 100 µm), *E*13 (scale bars, 250 µm), and *E*14 (scale bars, 250 µm) lungs from control and *Tbx4-rtTA; Tet-O-Cre; Wnt5a^fl/fl^* embryos. **E,** Normalized number of terminal branches in control and *Tbx4-rtTA; Tet-O-Cre; Wnt5a^fl/fl^* lungs at *E*12, *E*13 (*n*=5 control and *n*=7 mutant lungs, *p*=0.0351 via unpaired Student’s t-test), and *E*14 (*n*=6 control and *n*=7 mutant lungs, *p*=0.0478 via unpaired Student’s t-test). Different shapes represent different experimental replicates. Shown are mean ± s.d; * p < 0.05. **F-G,** Schematics of lung with landmarks used for quantification of bronchus length and diameter. **H-I,** Quantification of bronchus length or diameter vs. number of terminal branches in *E*12-14 cranial, medial, and accessory bronchi and L1 branch from control and *Tbx4-rtTA; Tet-O-Cre; Wnt5a^fl/fl^* lungs. Shown are best-fit curves and 95% confidence intervals.

**Supplemental Figure 6.**
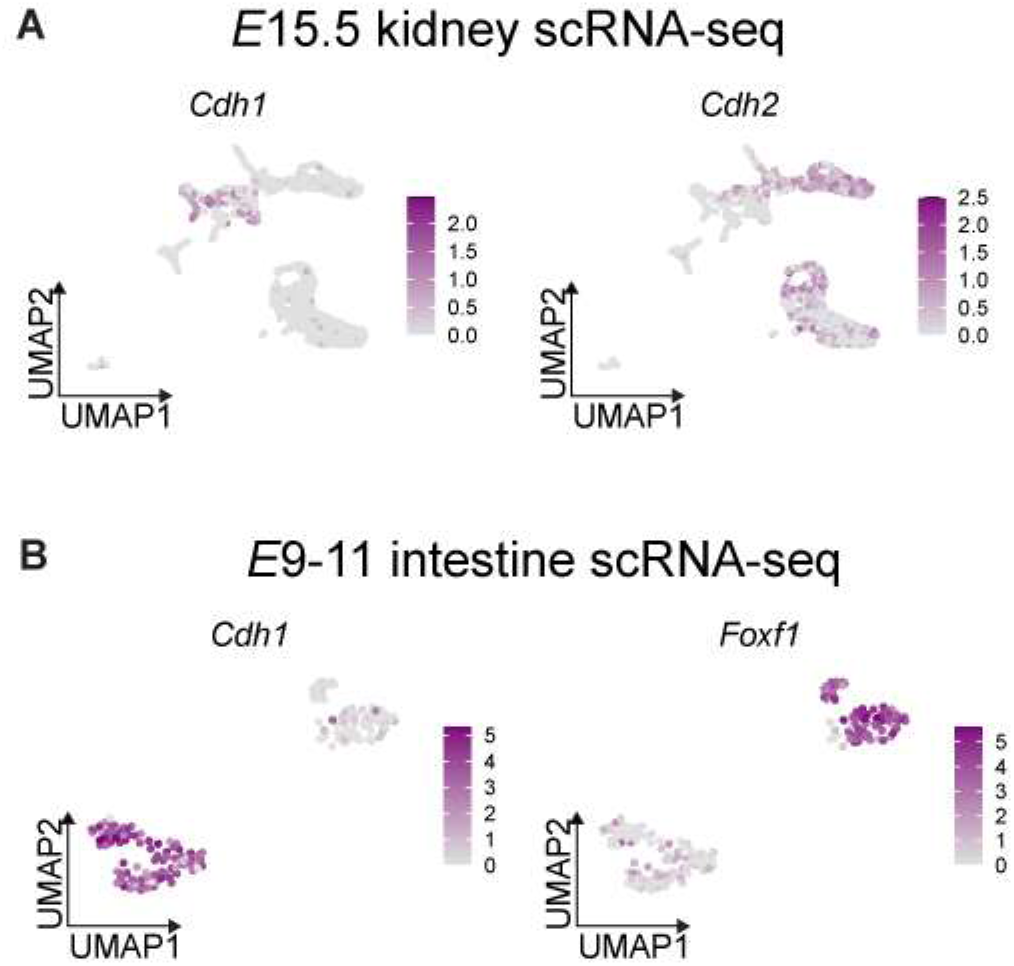
scRNAseq of developing organs. **A,** scRNAseq of developing kidney revealing epithelial (*Cdh1*-expressing) and mesenchymal (*Cdh2*-expressing) populations (Naganuma et al., 2021). **B,** scRNAseq of developing intestine revealing epithelial (*Cdh1*-expressing) and mesodermal (*Foxf1*-expressing) populations (Dong et al., 2018).

**Table 1.** Mouse Strains

| Mouse Line | Source or Reference | Strain |
| --- | --- | --- |
| *Vangl2<sup>Lp</sup>* | The Jackson Laboratory | JAX: 000220 |
| *Celsr1<sup>Crsh</sup>* | Elaine Fuchs, Jen Murdoch (Curtin et al., 2003) | MGI: 2668337 |
| *Shh<sup>tm1(EGFP/cre)Cjt</sup>* | The Jackson Laboratory | JAX: 005622 |
| *Dermo1Cre (Twist2<sup>tm1.1(cre)Dor</sup>)* | The Jackson Laboratory | JAX: 008712 |
| *Tbx4-rtTA* | Gift from Wei Shi (Zhang et al., 2013) | N/A |
| *tet-O-Cre* | The Jackson Laboratory | JAX: 006234 |
| *Vangl1 flox (Vangl1<sup>tm1.1Nat</sup>)* | The Jackson Laboratory | JAX: 019518 |
| *Vangl2 flox (Vangl2<sup>tm2.1Mdea</sup>)* | The Jackson Laboratory | JAX: 025174 |
| *Wnt5a flox (Wnt5a<sup>tm1.1Krvl</sup>)* | The Jackson Laboratory | JAX: 026626 |
| *mTmG reporter: (Gt(ROSA)26Sor<sup>tm4(ACTB-tdTomato,-EGFP)Luo</sup>)* | The Jackson Laboratory | JAX: 007676 |
| *Confetti reporter: (Gt(ROSA)26Sor<sup>tm1(CAG-Brainbow2.1)Cle</sup>)* | The Jackson Laboratory | JAX: 017492 |

**Table 2.**
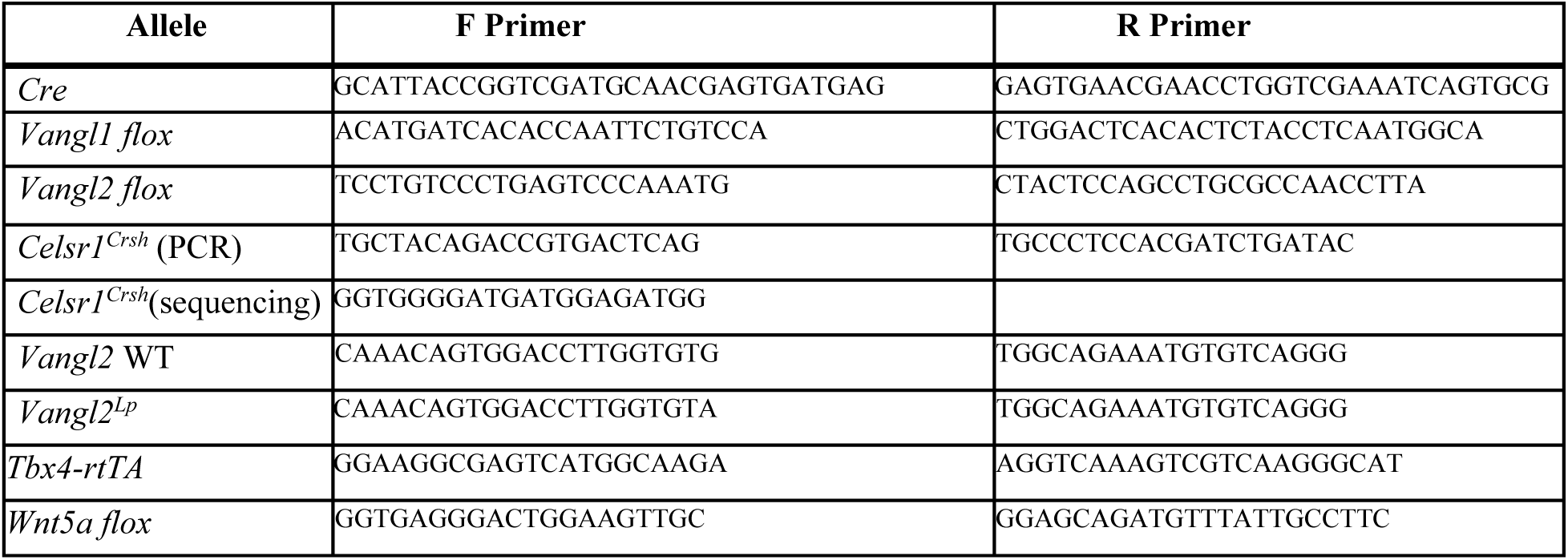
Genotyping Primers

**Table 3.** Antibodies

| **Antibody** | **Manufacturer, Cat#** | **Concentration** |
| --- | --- | --- |
| Ecad (rat) | Invitrogen, 14-3249-82 | 1:1000, frozen sections and whole mount |
| Ecad (rb) | Cell Signaling, 3195 | 1:200, frozen sections and whole mount |
| Celsr1 (guinea pig) | Custom Antibody (Danelle Devenport) | 1:1000, frozen sections and whole mount |
| Vangl2 (rat) | Millipore, MABN750 | 1:100, frozen sections and whole mount |
| Frizzled6 (goat) | R&D, AF1526 | 1:400, frozen sections and whole mount |
| GFP (chicken) | Abcam, ab13970 | 1:500, frozen sections |
| RFP, also recognizes tdTomato (rb) | Rockland Inc, #600-401-379 | 1:200, frozen sections |
| αSMA (mouse monoclonal) | Sigma, a5228 | 1:200, frozen sections and whole mount |
| Fibronectin (rb) | Custom Antibody (Jean Schwarzbauer) | 1:200, frozen sections |
| Collagen IV (rb) | Abcam, ab6586 | 1:200, frozen sections |
| Phospho-Histone-3 (rb) | Upstate (Millipore), 06-570 | 1:200, frozen sections |
| Cleaved Caspase-3 (rb) | Cell Signaling, #9661S | 1:200, frozen sections |
| Lef1 (rabbit monoclonal) | Cell Signaling, #2230 | 1:200, frozen sections |
| FoxP1 (rabbit polyclonal) | Cell Signaling, #2005 | 1:200, frozen sections |

## Notes

### Competing Interest Statement

The authors have declared no competing interest.

## References

1. Basta, L. P., Hill-Oliva, M., Paramore, S. V., Sharan, R., Goh, A., Biswas, A., Cortez, M., Little, K. A., Posfai, E. & Devenport, D. 2021. New mouse models for high resolution and live imaging of planar cell polarity proteins in vivo. Development, 148.

2. Boutin, C., Labedan, P., Dimidschstein, J., Richard, F., Cremer, H., Andre, P., Yang, Y., Montcouquiol, M., Goffinet, A. M. & Tissir, F. 2014. A dual role for planar cell polarity genes in ciliated cells. Proc Natl Acad Sci U S A, 111, E3129–38.

3. Butler, A., Hoffman, P., Smibert, P., Papalexi, E. & Satija, R. 2018. Integrating single-cell transcriptomic data across different conditions, technologies, and species. Nat Biotechnol, 36, 411–420.

4. Cetera, M., Leybova, L., Woo, F. W., Deans, M. & Devenport, D. 2017. Planar cell polarity-dependent and independent functions in the emergence of tissue-scale hair follicle patterns. Dev Biol, 428, 188–203.

5. Chen, H., Zhuang, F., Liu, Y. H., Xu, B., DEL Moral, P., Deng, W., Chai, Y., Kolb, M., Gauldie, J., Warburton, D., Moses, H. L. & Shi, W. 2008. TGF-beta receptor II in epithelia versus mesenchyme plays distinct roles in the developing lung. Eur Respir J, 32, 285–95.

6. Copley, C. O., Duncan, J. S., Liu, C., Cheng, H. & Deans, M. R. 2013. Postnatal refinement of auditory hair cell planar polarity deficits occurs in the absence of Vangl2. J Neurosci, 33, 14001–16.

7. Curtin, J. A., Quint, E., Tsipouri, V., Arkell, R. M., Cattanach, B., Copp, A. J., Henderson, D. J., Spurr, N., Stanier, P., Fisher, E. M., Nolan, P. M., Steel, K. P., Brown, S. D., Gray, I. C. & Murdoch, J. N. 2003. Mutation of Celsr1 disrupts planar polarity of inner ear hair cells and causes severe neural tube defects in the mouse. Curr Biol, 13, 1129–33.

8. Devenport, D. 2014. The cell biology of planar cell polarity. J Cell Biol, 207, 171–9.

9. Devenport, D. & Fuchs, E. 2008. Planar polarization in embryonic epidermis orchestrates global asymmetric morphogenesis of hair follicles. Nat Cell Biol, 10, 1257–68.

10. Dong, J., Hu, Y., Fan, X., Wu, X., Mao, Y., Hu, B., Guo, H., Wen, L. & Tang, F. 2018. Single-cell RNA-seq analysis unveils a prevalent epithelial/mesenchymal hybrid state during mouse organogenesis. Genome Biol, 19, 31.

11. Gao, B., Song, H., Bishop, K., Elliot, G., Garrett, L., English, M. A., Andre, P., Robinson, J., Sood, R., Minami, Y., Economides, A. N. & Yang, Y. 2011. Wnt signaling gradients establish planar cell polarity by inducing Vangl2 phosphorylation through Ror2. Dev Cell, 20, 163–76.

12. Gilbert, R. M., Schappell, L. E. & Gleghorn, J. P. 2021. Defective mesothelium and limited physical space are drivers of dysregulated lung development in a genetic model of congenital diaphragmatic hernia. Development, 148.

13. Goodwin, K., Jaslove, J. M., Tao, H., Zhu, M., Hopyan, S. & Nelson, C. M. 2020. Patterning the embryonic pulmonary mesenchyme. bioRxiv, 2020.08.20.259101.

14. Goodwin, K., Jaslove, J. M., Tao, H., Zhu, M., Hopyan, S. & Nelson, C. M. 2022. Patterning the embryonic pulmonary mesenchyme. iScience, 25, 103838.

15. Goodwin, K., Lemma, B., Zhang, P., Boukind, A. & Nelson, C. M. 2023. Plasticity in airway smooth muscle differentiation during mouse lung development. Dev Cell, 58, 338–347 e4.

16. Goodwin, K., Mao, S., Guyomar, T., Miller, E., Radisky, D. C., Kosmrlj, A. & Nelson, C. M. 2019. Smooth muscle differentiation shapes domain branches during mouse lung development. Development, 146, dev181172.

17. Goto, T. & Keller, R. 2002. The planar cell polarity gene strabismus regulates convergence and extension and neural fold closure in Xenopus. Dev Biol, 247, 165–81.

18. Harfe, B. D., Scherz, P. J., Nissim, S., Tian, H., Mcmahon, A. P. & Tabin, C. J. 2004. Evidence for an expansion-based temporal Shh gradient in specifying vertebrate digit identities. Cell, 118, 517–28.

19. Harris, K. S., Zhang, Z., Mcmanus, M. T., Harfe, B. D. & Sun, X. 2006. Dicer function is essential for lung epithelium morphogenesis. Proc Natl Acad Sci U S A, 103, 2208–13.

20. Heisenberg, C. P., Tada, M., Rauch, G. J., Saude, L., Concha, M. L., Geisler, R., Stemple, D. L., Smith, J. C. & Wilson, S. W. 2000. Silberblick/Wnt11 mediates convergent extension movements during zebrafish gastrulation. Nature, 405, 76–81.

21. Kibar, Z., Vogan, K. J., Groulx, N., Justice, M. J., Underhill, D. A. & Gros, P. 2001. Ltap, a mammalian homolog of Drosophila Strabismus/Van Gogh, is altered in the mouse neural tube mutant Loop-tail. Nat Genet, 28, 251–5.

22. Kim, H. Y., Pang, M. F., Varner, V. D., Kojima, L., Miller, E., Radisky, D. C. & Nelson, C. M. 2015. Localized Smooth Muscle Differentiation Is Essential for Epithelial Bifurcation during Branching Morphogenesis of the Mammalian Lung. Dev Cell, 34, 719–26.

23. Kishimoto, K., Tamura, M., Nishita, M., Minami, Y., Yamaoka, A., Abe, T., Shigeta, M. & Morimoto, M. 2018. Synchronized mesenchymal cell polarization and differentiation shape the formation of the murine trachea and esophagus. Nat Commun, 9, 2816.

24. Kunimoto, K., Bayly, R. D., Vladar, E. K., Vonderfecht, T., Gallagher, A. R. & Axelrod, J. D. 2017. Disruption of Core Planar Cell Polarity Signaling Regulates Renal Tubule Morphogenesis but Is Not Cystogenic. Curr Biol, 27, 3120–3131 e4.

25. Kunimoto, K., Weiner, A. T., Axelrod, J. D. & Vladar, E. K. 2022. Distinct overlapping functions for Prickle1 and Prickle2 in the polarization of the airway epithelium. Front Cell Dev Biol, 10, 976182.

26. Lienkamp, S. S., Liu, K., Karner, C. M., Carroll, T. J., Ronneberger, O., Wallingford, J. B. & Walz, G. 2012. Vertebrate kidney tubules elongate using a planar cell polarity-dependent, rosette-based mechanism of convergent extension. Nat Genet, 44, 1382–7.

27. Livet, J., Weissman, T. A., Kang, H., Draft, R. W., Lu, J., Bennis, R. A., Sanes, J. R. & Lichtman, J. W. 2007. Transgenic strategies for combinatorial expression of fluorescent proteins in the nervous system. Nature, 450, 56–62.

28. Metzger, R. J., Klein, O. D., Martin, G. R. & Krasnow, M. A. 2008. The branching programme of mouse lung development. Nature, 453, 745–50.

29. Naganuma, H., Miike, K., Ohmori, T., Tanigawa, S., Ichikawa, T., Yamane, M., Eto, M., Niwa, H., Kobayashi, A. & Nishinakamura, R. 2021. Molecular detection of maturation stages in the developing kidney. Dev Biol, 470, 62–73.

30. Nelson, C. M., Gleghorn, J. P., Pang, M. F., Jaslove, J. M., Goodwin, K., Varner, V. D., Miller, E., Radisky, D. C. & Stone, H. A. 2017. Microfluidic chest cavities reveal that transmural pressure controls the rate of lung development. Development, 144, 4328–4335.

31. Nelson, T. R., West, B. J. & Goldberger, A. L. 1990. The fractal lung: universal and species-related scaling patterns. Experientia, 46, 251–4.

32. Paramore, S. V., Trenado-Yuste, C., Sharan, R., Devenport, D. & Nelson, C. M. 2022. Vangl facilitates mesenchymal thinning during lung sacculation independently of Celsr. bioRxiv, 2022.12.28.522148.

33. Paudyal, A., Damrau, C., Patterson, V. L., Ermakov, A., Formstone, C., Lalanne, Z., Wells, S., Lu, X., Norris, D. P., Dean, C. H., Henderson, D. J. & Murdoch, J. N. 2010. The novel mouse mutant, chuzhoi, has disruption of Ptk7 protein and exhibits defects in neural tube, heart and lung development and abnormal planar cell polarity in the ear. BMC Dev Biol, 10, 87.

34. Rao-Bhatia, A., Zhu, M., Yin, W. C., Coquenlorge, S., Zhang, X., Woo, J., Sun, Y., Dean, C. H., Liu, A., Hui, C. C., Shivdasani, R. A., Mcneill, H., Hopyan, S. & Kim, T. H. 2020. Hedgehog-Activated Fat4 and PCP Pathways Mediate Mesenchymal Cell Clustering and Villus Formation in Gut Development. Dev Cell, 52, 647–658 e6.

35. Ryu, Y. K., Collins, S. E., Ho, H. Y., Zhao, H. & Kuruvilla, R. 2013. An autocrine Wnt5a-Ror signaling loop mediates sympathetic target innervation. Dev Biol, 377, 79–89.

36. Smith, P., Godde, N., Rubio, S., Tekeste, M., Vladar, E. K., Axelrod, J. D., Henderson, D. J., Milgrom-Hoffman, M., Humbert, P. O. & Hinck, L. 2019. VANGL2 regulates luminal epithelial organization and cell turnover in the mammary gland. Sci Rep, 9, 7079.

37. Sosic, D., Richardson, J. A., Yu, K., Ornitz, D. M. & Olson, E. N. 2003. Twist regulates cytokine gene expression through a negative feedback loop that represses NF-kappaB activity. Cell, 112, 169–80.

38. Stahley, S. N., Basta, L. P., Sharan, R. & Devenport, D. 2021. Celsr1 adhesive interactions mediate the asymmetric organization of planar polarity complexes. Elife, 10, e62097.

39. Sutherland, A., Keller, R. & Lesko, A. 2020. Convergent extension in mammalian morphogenesis. Semin Cell Dev Biol, 100, 199–211.

40. Tada, M. & Smith, J. C. 2000. Xwnt11 is a target of Xenopus Brachyury: regulation of gastrulation movements via Dishevelled, but not through the canonical Wnt pathway. Development, 127, 2227–38.

41. Tanabe, N., Sato, S., Suki, B. & Hirai, T. 2020. Fractal Analysis of Lung Structure in Chronic Obstructive Pulmonary Disease. Front Physiol, 11, 603197.

42. Tang, N., Marshall, W. F., Mcmahon, M., Metzger, R. J. & Martin, G. R. 2011. Control of mitotic spindle angle by the RAS-regulated ERK1/2 pathway determines lung tube shape. Science, 333, 342–345.

43. Tang, Z., Hu, Y., Wang, Z., Jiang, K., Zhan, C., Marshall, W. F. & Tang, N. 2018. Mechanical Forces Program the Orientation of Cell Division during Airway Tube Morphogenesis. Dev Cell, 44, 313–325 e5.

44. Torban, E., Wang, H. J., Groulx, N. & Gros, P. 2004. Independent mutations in mouse Vangl2 that cause neural tube defects in looptail mice impair interaction with members of the Dishevelled family. J Biol Chem, 279, 52703–13.

45. Vladar, E. K., Bayly, R. D., Sangoram, A. M., Scott, M. P. & Axelrod, J. D. 2012. Microtubules enable the planar cell polarity of airway cilia. Curr Biol, 22, 2203–12.

46. Vladar, E. K., Nayak, J. V., Milla, C. E. & Axelrod, J. D. 2016. Airway epithelial homeostasis and planar cell polarity signaling depend on multiciliated cell differentiation. JCI Insight, 1.

47. Wallingford, J. B., Rowning, B. A., Vogeli, K. M., Rothbacher, U., Fraser, S. E. & Harland, R. M. 2000. Dishevelled controls cell polarity during Xenopus gastrulation. Nature, 405, 81–5.

48. Wang, Y., Guo, N. & Nathans, J. 2006. The role of Frizzled3 and Frizzled6 in neural tube closure and in the planar polarity of inner-ear sensory hair cells. J Neurosci, 26, 2147–56.

49. Yates, L. L., Schnatwinkel, C., Murdoch, J. N., Bogani, D., Formstone, C. J., Townsend, S., Greenfield, A., Niswander, L. A. & Dean, C. H. 2010. The PCP genes Celsr1 and Vangl2 are required for normal lung branching morphogenesis. Hum Mol Genet, 19, 2251–67.

50. Yin, H., Copley, C. O., Goodrich, L. V. & Deans, M. R. 2012. Comparison of phenotypes between different vangl2 mutants demonstrates dominant effects of the Looptail mutation during hair cell development. PLoS One, 7, e31988.

51. Yin, W., Kim, H. T., Wang, S., Gunawan, F., Wang, L., Kishimoto, K., Zhong, H., Roman, D., Preussner, J., Guenther, S., Graef, V., Buettner, C., Grohmann, B., Looso, M., Morimoto, M., Mardon, G., Offermanns, S. & Stainier, D. Y. R. 2018. The potassium channel KCNJ13 is essential for smooth muscle cytoskeletal organization during mouse tracheal tubulogenesis. Nat Commun, 9, 2815.

52. Yin, Y., White, A. C., Huh, S. H., Hilton, M. J., Kanazawa, H., Long, F. & Ornitz, D. M. 2008. An FGF-WNT gene regulatory network controls lung mesenchyme development. Dev Biol, 319, 426–36.

53. Yu, K., Xu, J., Liu, Z., Sosic, D., Shao, J., Olson, E. N., Towler, D. A. & Ornitz, D. M. 2003. Conditional inactivation of FGF receptor 2 reveals an essential role for FGF signaling in the regulation of osteoblast function and bone growth. Development, 130, 3063–74.

54. Yuan, T., Volckaert, T., Chanda, D., Thannickal, V. J. & De Langhe, S. P. 2018. Fgf10 Signaling in Lung Development, Homeostasis, Disease, and Repair After Injury. Front Genet, 9, 418.

55. Zhang, K., Yao, E., Chuang, E., Chen, B., Chuang, E. Y., Volk, R. F., Hofmann, K. L., Zaro, B. & Chuang, P. T. 2022. Wnt5a-Vangl1/2 signaling regulates the position and direction of lung branching through the cytoskeleton and focal adhesions. PLoS Biol, 20, e3001759.

56. Zhang, K., Yao, E., Lin, C., Chou, Y. T., Wong, J., Li, J., Wolters, P. J. & Chuang, P. T. 2020. A mammalian Wnt5a-Ror2-Vangl2 axis controls the cytoskeleton and confers cellular properties required for alveologenesis. Elife, 9, e53688.

57. Zhang, W., Menke, D. B., Jiang, M., Chen, H., Warburton, D., Turcatel, G., Lu, C. H., Xu, W., Luo, Y. & Shi, W. 2013. Spatial-temporal targeting of lung-specific mesenchyme by a Tbx4 enhancer. BMC Biol, 11, 111.

